# Structure of a key adhesin-exopolysaccharide interaction provides insights into matrix assembly in *Vibrio cholerae* biofilms

**DOI:** 10.64898/2026.08.03.742583

**Authors:** Alexander J. Hinbest, Hyerim Bianca Nam, Ewelina D. Liszczyk, Rajan Kandel, Emma Gerace, Alexis Moreau, Ranjuna Weerasekera, Ryan Gordon, Yiyan Yang, Jack Chen, Nathan Fowler, Xiaofang Jiang, Robert J. Woods, Jing Yan, Rich Olson

## Abstract

Biofilms serve as a protective mechanism for bacteria, including many pathogens. To form such communities, bacteria secrete macromolecules that form an extracellular matrix serving as a barrier against environmental threats, such as predation, antibiotics, and the host immune system. To be effective, a biofilm must anchor to foreign surfaces and retain sufficient stiffness, in the environment or in a host. However, how this matrix self-organizes to support biofilm formation remains a mystery at the molecular level. The human pathogen *Vibrio cholerae* produces biofilms primarily composed of an exopolysaccharide called VPS (<u>V</u>ibrio <u>p</u>oly<u>s</u>accharide), consisting of an unusually-modified repeating tetrasaccharide core unit. VPS engages with two secreted adhesion proteins, Bap1 and RbmC, which adhere the biofilm to abiotic and biotic surfaces, and serve to strengthen the biofilm by interacting with VPS using a conserved β-propeller. To pinpoint the interaction between the adhesins and purified segments of VPS, we determined the ∼1.6 Å X-ray crystal structure of Bap1 bound to fragmented VPS and used the structure to carry out molecular dynamics simulations. The structure revealed a single binding site consisting of one tetrasaccharide unit involving an induced magnesium binding site. Unexpectedly, the tetrasaccharide adopted a bent state caused by a rotation of the glycosidic bond between the central two monosaccharide units. Using a combination of mutagenesis, light scattering, and *in situ* fluorescent microscopy, we demonstrate that Bap1 not only facilitates biofilm adhesion, but is also required for proper VPS organization, through the identified binding pocket. Our structure reveals for the first time the interaction between a biofilm exopolysaccharide and matrix protein, as well as insights into conformational changes of exopolysaccharide induced by this binding. Our findings provide a generalizable approach for studying the biophysical and biochemical properties of carbohydrate-dependent biofilm assembly, which may lead to new ways to treat disease caused by biofilm-forming bacterial pathogens by disrupting the exopolysaccharide-protein interactions.

## Introduction

To protect themselves from environmental threats, most bacteria form biofilms, which consist of a matrix of secreted extracellular factors held together via molecular interactions^1^. In addition to forming a protective sheath around growing bacterial colonies, the extracellular matrix also enables the biofilm-dwelling cells to adhere to various surfaces encountered in their lifecycle^2^. This may be abiotic surfaces in the environment, or in the case of symbiotic or pathogenic species, specific receptors found on a host organism. While some bacteria use direct surface-anchored adhesins to achieve host adhesion, the majority of biofilm-forming species utilize secreted factors including exopolysaccharides and matrix proteins^3^. Hence, in addition to the attachment to foreign surfaces, these secreted factors may also interact with each other and exopolysaccharides in the matrix. This recognition often takes place in the presence of an overwhelming amount of sugar moieties from the host in the form of *O*-or *N*-glycans^4^, which can potentially interfere with the binding of matrix proteins and exopolysaccharides. Very little is known about the molecular details underlying these interactions, which hinders our understanding of the fundamental principles underlying the assembly of the biofilm matrix. Understanding how matrix components combine at the molecular level to protect bacteria and allow them to colonize hosts is essential in the fight against biofilm-related diseases and particularly important in the age of growing bacterial persistence and resistance to antibiotic treatments^5,6^.

*Vibrio cholerae* is an aquatic bacterium that causes the severe human diarrheal disease cholera, leading to hundreds of thousands of cases and thousands of deaths worldwide, annually^7,8^. Biofilm formation has been implicated to play significant roles during its perseverance in nature and infection in the host^9^. The biofilm matrix of *V. cholerae* consists of a secreted polysaccharide called *<u>V</u>ibrio* <u>p</u>oly<u>s</u>accharide (VPS), as well as several extracellular matrix proteins that serve to glue biofilm cells together and to surfaces^9–11^. The structure of VPS has been determined to consist of a repeating tetrasaccharide with the following chemical makeup: α-L-GulNAcAGly3OAc-(1-4)-β-D-Glc-(1-4)-α-D-Glc-(1-4)-α-D-Gal-(1-4)-^12^ (Fig. 1A). The L-gulose moiety found within the polymer is uniquely modified by an N-acetyl group at C2, an O-acetyl group at C3, and an amide-linked glycine residue at C6, imparting one negative charge per repeating unit. A recent crystal structure of the *V. cholerae* biofilm dispersal enzyme RbmB bound to tetra-and octa-saccharide fragments of the VPS polymer suggested that the glycine modification pays a role in the recognition of the polymer through salt-bridging interactions with arginine residues found within the VPS binding cleft^13^.

**Figure 1.**
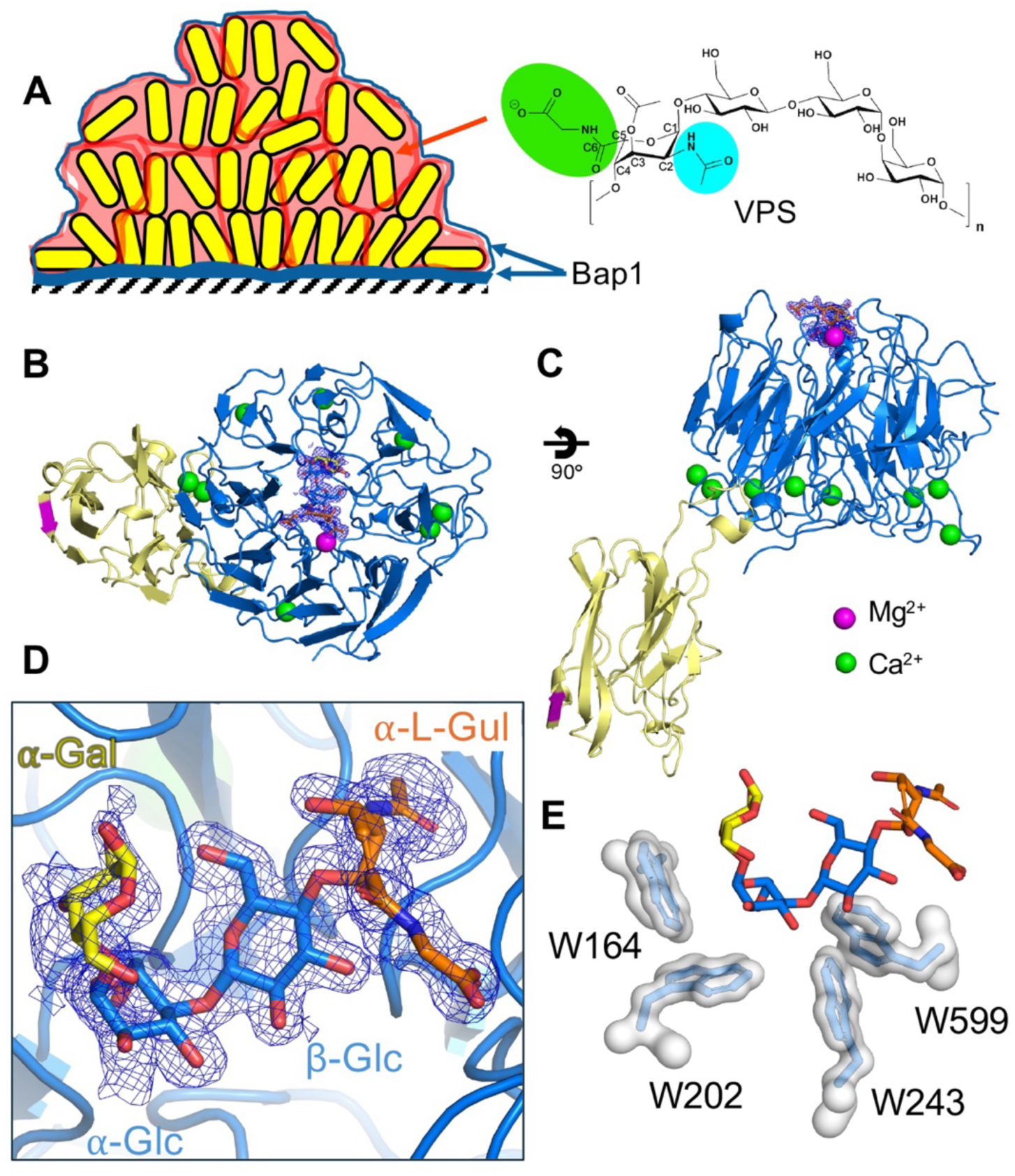
Crystal structure of Bap1-VPS complex. (A) Schematic showing functional location of Bap1 adhesin (blue line) and VPS (red lines) throughout biofilm. Bap1 bridges the biofilm and substrate, binding VPS on one side and the surface on the other; it additionally colocalizes with VPS surrounding the biofilm. Structure of the VPS repeating unit is shown to the right with amide-linked glycine and N-acetyl group highlighted with green and cyan ovals, respectively. (B) Top and side (C) views of structural model for Bap1-VPS showing Poulder map density contoured to 1σ in blue mesh. The VPS-binding β-propeller domain is in blue, the β-prism in yellow, and the location of the removed 57-amino acid adhesion loop that binds to surfaces is in magenta. Magenta and green spheres indicate Mg^2+^ and Ca^2+^ sites, respectively. (D) Closeup of VPS binding site showing the tetramer in stick conformation. Sugars are colored using Symbol Nomenclature for Glycans (SNFG) representation^72^. (E) Four tryptophan residues cradle VPS in the unusual bent conformation.

In addition to VPS, *V. cholerae* secretes several matrix proteins (RbmA, RbmC, Bap1)^14^ known to interact with the exopolysaccharide in different ways. RbmA^15^, which consists of a dimeric fibronectin type III fold^16,17^, binds VPS^18^ and has been proposed to interact with *V. cholerae* cell surfaces^19,20^ to facilitate cell-cell adhesion. In addition, *V. cholerae* also secretes two adhesion proteins, Bap1 and RbmC, with partially redundant functions^14^, that are the primary determinants of biofilm adhesion to foreign surfaces^2,20,21^. The high-resolution structure of Bap1 has been determined to high resolution revealing an 8-bladed β-propeller domain^21^ and a β-prism domain with structural similarities to plant lectin domains^22^. The β-prism domain in Bap1 does not appear to exhibit sugar-binding activity, but rather contains an intervening 57-amino acid peptide loop with similarities to mussel foot proteins that facilitates binding to abiotic surfaces^21^ and membranes^23^. RbmC shares the β-propeller domain with Bap1, and additionally contains two lectin domains with β-prism folds that bind with nanomolar affinity to the core branch of *N*-linked glycans found ubiquitously on animal cell surfaces^24^. RbmC also contains two additional βγ-crystallin domains that bind mucins^21,25^, another common component found on animal epithelial surfaces like the human gut. Together, Bap1 and RbmC allow *V. cholerae* biofilms to interact with a wide variety of abiotic and biotic surfaces, and may contribute to the efficacy of the bacterium to colonize the human gut^26^.

Although Bap1 and RbmC contain diverse accessory domains to facilitate binding to various environmental surfaces, they share a β-propeller domain with approximately 70% sequence identity^14^. We previously demonstrated that the β-propeller domain directly binds VPS, and this binding leads to the formation of large aggregates^27^. Unfortunately, it is not possible to study the interaction between β-propeller domains and VPS using crystallography due to the size, flexibility, and heterogeneity of VPS polymers, necessitating a different approach. We took advantage of a recent discovery that another protein found within the cluster of biofilm-related genes in *V. cholerae* and hypothesized to be a dispersal factor^14,28^, RbmB, acts as a polysaccharide lyase to cleave VPS into fragments^13,27^ that are more conducive to a crystallographic approach.

Here, we co-crystallized Bap1 with a mixture of cleaved fragments of VPS and determined the structure using X-ray diffraction. We report the structure of the Bap1-VPS complex at approximately 1.6 Å resolution. Our structure revealed clear electron density for a tetrasaccharide unit of VPS bound to a site at the center of the β-propeller domain, providing key insights into the mechanism of VPS recognition. Our structure, associated functional experiments, and molecular dynamics (MD) simulations provide a model for understanding how binding of VPS by Bap1 may lead to aggregation and consequently strengthening the biofilm matrix. Along with the previously determined structure of RbmB bound to VPS^13^, our structure reveals details about how matrix proteins with completely different folds uniquely recognize the same polymer utilizing the unique chemical modifications found on the L-gulose unit within the VPS tetramer. Our investigation provides insights into the mechanism of *V. cholerae* biofilm assembly and adhesion, as well as serving as a template for studying exopolysaccharide-protein interactions in biofilms from other bacterial systems that have evaded structural investigation so far.

## Results

### Crystal structure determination of Bap1 bound to VPS tetrasaccharide

To determine the structure of Bap1 bound to VPS fragments, we utilized the polysaccharide lyase RbmB from *V. cholerae* to cleave purified VPS into fragments. Due to the extended binding footprint of RbmB^13^, the enzyme produces a mixture of tetrasaccharide and octasaccharide fragments of VPS by cleaving the α(1-4) bond between the D-galactose and modified L-gulose monosaccharides^27^. Acting in a typical polysaccharide lyase β-elimination reaction^29^, this cleavage also modifies the resulting L-gulose moiety, leaving an unsaturated C-C double bond between C4 and C5, but leaving the rest of the cleaved fragments chemically intact. Concentrated VPS (∼0.7mM) was co-crystallized with Bap1_Δ57_-a construct missing the 57-amino acid adhesive loop (required for solubility, referred to as just Bap1 hereafter) yielding crystals with a similar morphology and space group as previously observed for the Apo structure^22^. The structure was solved and refined as described in Methods and Materials and the final model was refined to an R_work_ and R_free_ of 15.4, and 17.6, respectively (Tables S1 and S2).

During the process of refinement, additional electron density was apparent within a pocket at the center of the Bap1 β-propeller domain, which clearly resembled an oligosaccharide (Fig. 1B,C). Interpretation of the electron density allowed us to model a single tetrasaccharide unit of VPS, starting with the modified L-gulose and ending with the terminal α-galactose. Annealed F_o_-F_c_ Polder maps generated by phenix.refine^30^ indicated excellent electron density for the first three monosaccharides, while the α-galactose moiety exhibited less-well defined electron density and higher B-factors suggesting heterogeneity or thermal motion (Fig. 1D and Fig. S1). We see clear electron density for the N-acetyl and glycine groups on the modified L-gulose sugar, but do not observe density for the O-acetyl group, which we hypothesize may be chemically labile and lost during the VPS purification and cleavage procedure. Notably, there is ample room around the C3 hydroxyl group to accommodate such a group if it were present.

It has also been determined that ∼20% of the α-D-Glc sugar may be replaced by N-acetylglucosamine (GlcNAc)^12^. We observe both smeared density in the vicinity of the C2 position in the α-D-Glc sugar and ample room to accommodate the group, the density was not well-defined enough to justify building in a partially occupied GlcNAc. Although we expect that octasaccharide fragments were present in the crystallization mixture^13,27^, we did not see any evidence in the maps for such, and we also did not find any extra electron density within the protein structure consistent with additional bound VPS chains.

### Structural analysis of the VPS binding pocket

The bound VPS tetramer sits close to the center of the 8-fold pseudosymmetry axis of the 8-bladed β-propeller domain (Fig. 1B), adopting an unusually bent “horseshoe-like” conformation cradled by four tryptophan residues (W164, W202, W243, and W599 – Fig. 1E). Aromatic-carbohydrate interactions are a hallmark of many lectins, where CH-π interactions contribute to the overall binding energy and positioning of carbohydrate ligands within binding sites^31^. A closer examination of the individual tryptophan residues indicated a range of protein/carbohydrate configurations involved in each specific interaction (Fig. S2). Whereas W164 interacts in a parallel fashion with the hydrophobic face of the α1,4 glycosidic bond between the α-Glc C1 and α-Gal C4 positions, W202 interacts with the α-Glc ring at a roughly 45 degrees angle (Fig. S2).

The final two residues, W243 and W599, interact perpendicularly to the β1,4 glycosidic bond and β-Glc ring, respectively.

To analyze the specific interactions that anchor the VPS molecule into the Bap1 binding site, we used the program LigPlot+ (v. 2.2.8)^32^ to identify and map polar and non-polar interactions (Fig. 2A). In refining the structure, we also noticed a sphere of electron density interacting with the N-acetyl group on the L-gulose sugar that was too highly coordinated to suggest a water molecule. Analysis of the model using the CheckMyMetal algorithm (https://cmm.minorlab.org/) suggested magnesium, manganese, or cobalt as the most likely ions based on an octahedral geometry and bond lengths around 2 Å. We modeled a Mg^2+^ ion in this position as it is the most likely to have been present during purification and crystallization. The ion is coordinated by an unusual backbone conformation around T597 leading to a Ramachandran outlier of this residue. The backbone carbonyls of G596 and T597 along with the sidechain hydroxyl of T597 make up half of the coordination geometry, with the sidechain hydroxyl of S551 and sidechain carbonyl of N550 completing the site. Superposition of the area around the ion binding site with the Apo structure (PDB:6MLT) illustrate structural rearrangements in residues N550 and S551 that accompany coordination of the ion (Fig. 2B). As we observe no metal ion bound here in the Apo Bap1 structure, this suggests that binding of the VPS tetrasaccharide leads to the formation of this ion binding site. Indeed, in the aquatic and host environment where *V. cholerae* biofilms form, many ions including magnesium would be abundant.

**Figure 2.**
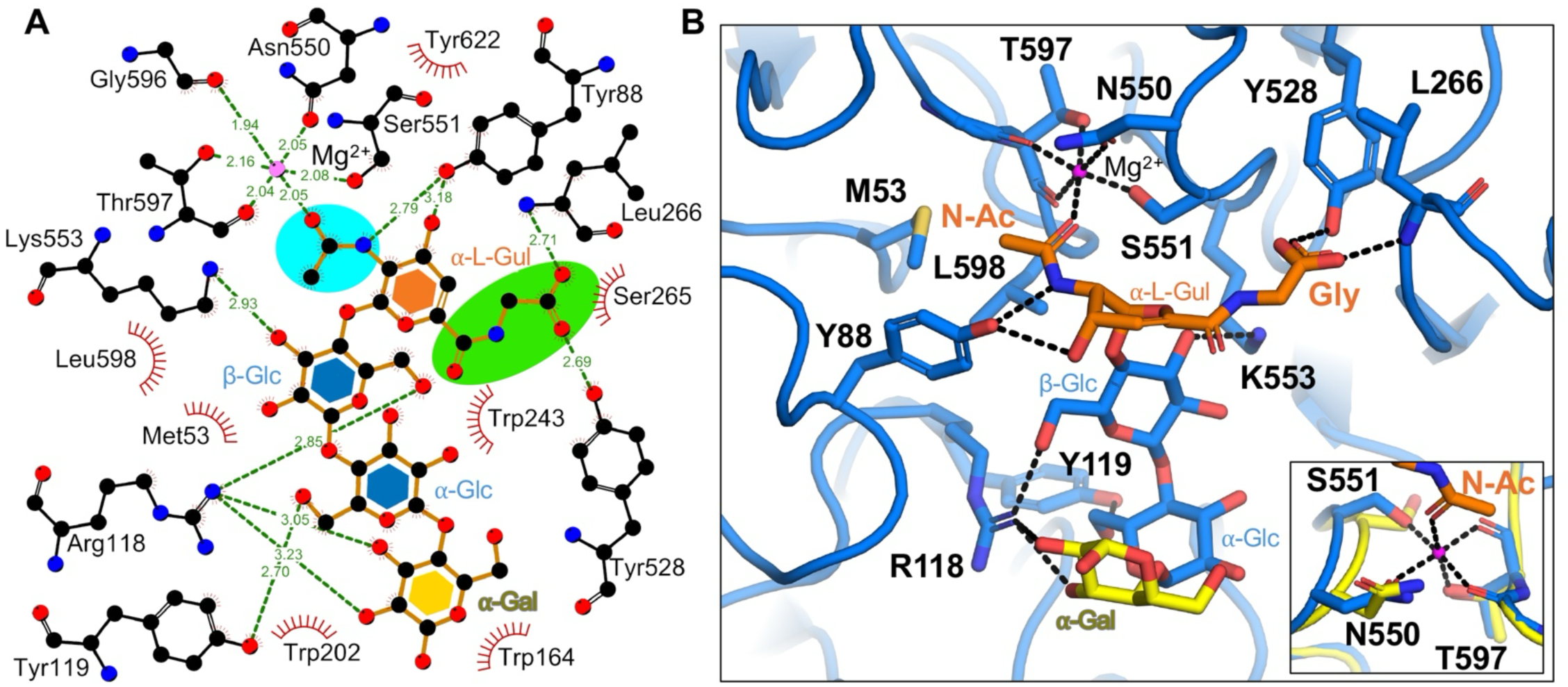
Structural details of VPS binding site. (A) Ligplot+ schematic outlining interactions between Bap1 and VPS. Hydrogen bonding interactions with distances in Å are shown as green dotted lines and van der Waals interactions as red crescents. A single Mg^2+^ ion is shown as a magenta sphere. Amide-linked glycine and N-acetyl groups are highlighted with green and cyan ovals, respectively. (B) Three-dimensional image of Bap1-VPS interactions with hydrogen bonds denoted by black dotted lines. Bottom-right inset, a superposition of ion-coordinating amino acid residues for Apo (yellow, PDB:6MLT) and VPS-bound structures (blue) illustrates conformational changes that accompany ion binding.

The same VPS N-acetyl group that interacts with the bound ion also projects its methyl group into a hydrophobic pocket formed by M53, Y88, and L598 (Fig. 2B). The VPS amide linked glycine moiety forms putative hydrogen bond interactions with the sidechain hydroxyl of Y528 as well as the backbone amine group of L266. Since the VPS tetramer in this crystal structure was produced by a glycoside lyase, a distorted heterocyclic ring with an unsaturated bond between C4 and C5 is observed. It is unlikely that Bap1 would be interacting with heterocyclic gulose moiety under physiological conditions, and so it is likely that the model represents a slightly distorted version of native interactions. Nevertheless, we do not see any possible salt-bridge forming residues within the vicinity of the glycine moiety as was seen interacting with this group in two out of the three VPS repeating units in the RbmB crystal structure^13^.

The rest of the polar interactions between Bap1 and VPS mostly consisted of hydrogen-bonding interactions. This includes the sidechain hydroxyl group of Y88, which makes putative hydrogen-bonds with the nitrogen in the L-gulose N-acetyl group and the C3 hydroxyl on the L-gulose ring. Additional hydrogen bonds are formed between VPS glycan hydroxyl groups and the sidechains of R118, Y119, and K553, (Fig. 2A,B). In total, the VPS ligand buried approximately 568 Å^2^, or 480 Å^2^, of solvent-accessible surface-area as calculated by PISA (v. 1.52)^33^, or PyMOL (v. 3.1.6.1)^34^, respectively.

### VPS bending by Bap1

Due to the general rigidity of the monosaccharide heterocyclic rings, the configuration of a polysaccharide chain is mostly determined by the dihedral angles, Φ and Ψ, around the glycosidic bond linking sugars (measured as Φ: O_5_-C_1_-O_4_’-C_4_’ and Ψ: C_1_-O_4_’-C_4_’-C_3_’). To analyze the horseshoe-like bend in the VPS tetramer, we measured these angles for the three glycosidic bonds present in the crystal structure and compared them to the idealized GLYCAM-Web model (Table S3). We also compared them to the VPS tetramer found within the RbmB crystal structure (PDB: 9OZD), which adopts a linear conformation. We found that while the RbmB-bound VPS tetramer resembled the dihedral angles for the GLYCAM-Web model (all angles within 10° of each other), the Bap1-bound VPS tetramer dihedral angles diverged far from the other two models (Table S3). In particular, the central D-Glcβ(1-4)-D-Glcpα exhibited a roughly 170° rotation around the Ψ angle of the connecting glycosidic bond as compared to both the GLYCAM-Web and RbmB models. This result indicates that the bend observed in the modeled Bap1-VPS polymer results primarily from a rotation around the Glc-Glc glycosidic bond.

### Molecular dynamics simulations of VPS and Bap1-VPS

To further study the Bap1-VPS structure and the significance of the unusual D-Glcβ(1-4)-D-Glcα dihedral angle, we conducted MD simulations on a tetrasaccharide model initiated from the crystal structure and a dodecasaccharide (dodecamer) VPS model (consisting of three VPS tetramers). In both cases, the unsaturated bond within the L-gulose sugar was removed and the ring was modified to a typical chair conformation so that the presumed physiological state of the VPS polymer would be simulated. Triplicate independent MD simulations were performed for the Bap1 complexes with the tetramer and dodecamer VPS fragments (for 500 ns). Throughout all simulations, the VPS remained stably bound to Bap1, with crystallographically-observed hydrogen bonds present throughout the MD simulations (Table S4). Higher mobility was observed near the terminal glycan moieties of the VPS dodecamer that were not in direct contact with the protein (Fig. 3A, Fig. S3 and Table S5). Theoretical per-residue interaction energies were computed for each complex, which identified several key residues (Fig. S4 and Tables S6-S7). This includes residues R118 and L598, which appear to be key residues in stabilizing VPS binding. The most significant residue identified in the MD analysis was W164, which interacts with the somewhat flexible galactose residue in the crystal structure. Based on the per-residue interaction energy analysis, the modified L-gulose sugar contributed more than the other VPS residues towards Bap1 binding (Fig. S5 and Table S8).

**Figure 3.**
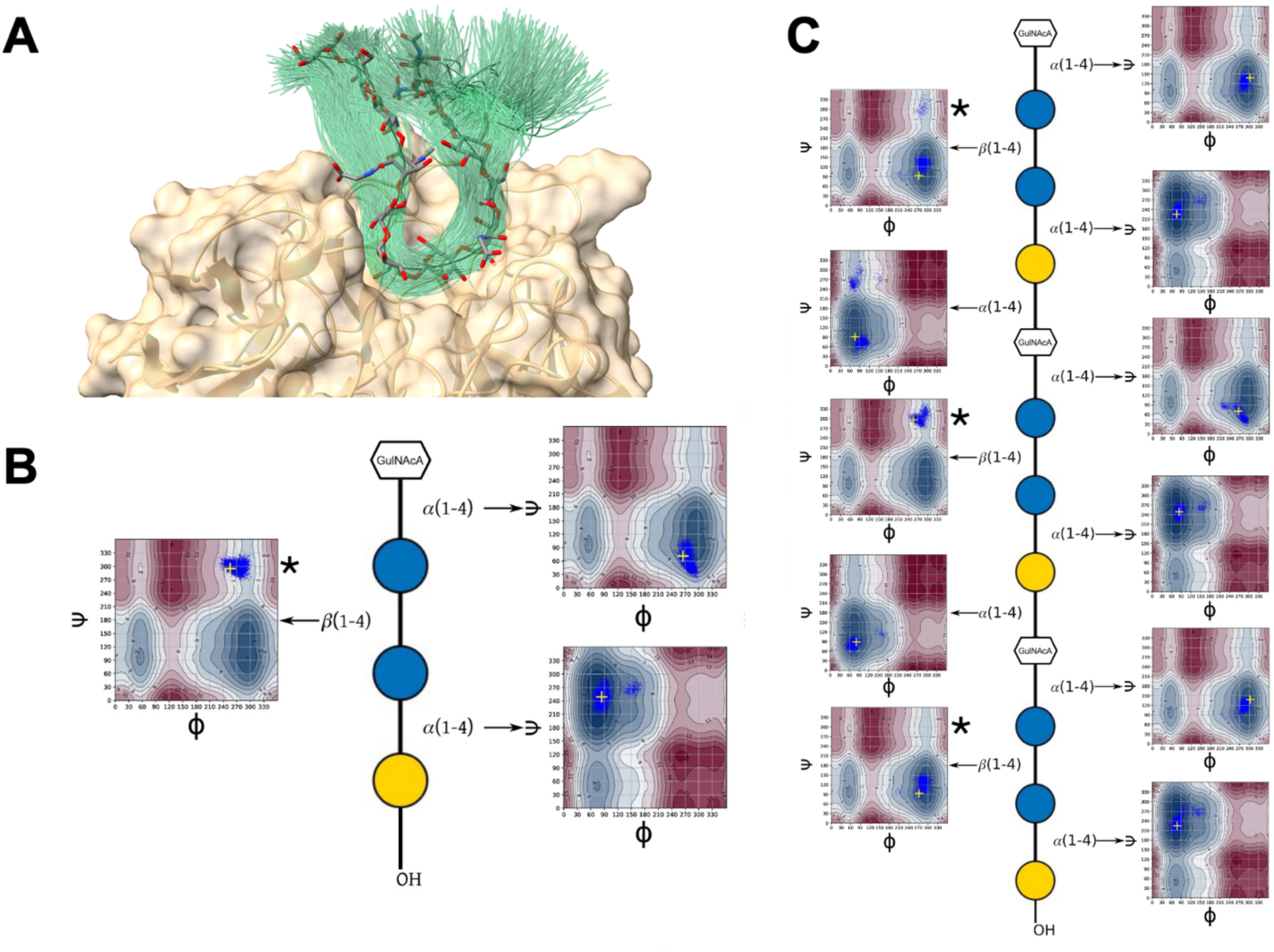
Molecular dynamics simulations of Bap1-VPS complex. (A) Representation of the motion of the bound dodecasaccharide model, indicated as a smoothed trace line (green) through the geometric centroids of each monosaccharide, shown for 500 frames selected at 1 ns timesteps from the 500 ns MD simulation. The single structure most similar to the average oligosaccharide shape is shown in licorice bond format. (B) Glycosidic angle distributions (Φ: O5-C1-O4-C4 and Ψ: C1-O4-C4-C3) from MD simulations of tetra-saccharide substrate fragments in complex with Bap1. (C) A model of the intact dodecasaccharide substrate in complex with Bap1. Contour plots indicate the theoretical glycosidic CHI energies. Starting conformations are indicated by a cross and black asterisks mark high energy Glc-Glc bond regions.

To better understand the energy landscape of glycosidic bonds within the VPS polymer, we plotted the carbohydrate intrinsic (CHI) energies for the oligosaccharide linkages (Fig. 3B,C). For the VPS tetrasaccharide (tetramer), the modified Gul-Glc and Glc-Gal dihedral angles remained within the low energy regions of the CHI-energy throughout the simulation (Fig. 3B). Removing the Mg^2+^ ion from the starting model led to unbinding of the VPS tetramer in two of three trials, emphasizing the importance of this feature for stabilizing the bound VPS molecule (Fig. S6). The Glc-Glc bond in the bound VPS tetramer also remained stable throughout the simulation, but occupied the higher energy anti-Ψ or “flipped” state observed in the crystal structure. The unusual nature of this feature is highlighted in the VPS dodecamer simulation, in which the central bound tetramer motif remained in this flipped conformation, while the two flanking tetramers, which do not directly contact Bap1, largely occupy the lower energy state seen in simulations of unbound VPS^13^ (Fig. 3C). These results suggest that it is binding with Bap1 that induces the high-energy conformation of the Glc-Glc bond in bound VPS.

### In vitro mutagenesis of VPS binding site

To better understand the individual contributions to the various interactions observed between Bap1 and VPS, we introduced point mutations in Bap1, and assessed their impact on Bap1-VPS binding. To start, we employed an assay to assess the ability of Bap1 mutants (using a Bap1-GFP_UV_ fusion protein) to bind to VPS in a native gel-shift experiment. When incubated with polymeric VPS, Bap1 shifts to a higher apparent molecular mass (barely entering the gel) as described previously^21^. To characterize relative binding, we started with a ratio of wild-type (WT) protein that led to a complete shift of the Bap1 signal to the higher molecular weight. We then assessed the amount of each Bap1 mutant that did not shift, which gave a semi-quantitative insight into the relative disruption of VPS binding of each mutant and allowed us to assess the relative impact of Bap1 binding site mutations on the VPS interaction (Fig. 4A and Fig. S7). We calculated the normalized unbound fraction of each Bap1 mutant, with a value of 1.0 being similar to WT (completely bound) and 0 representing no binding.

**Figure 4.**
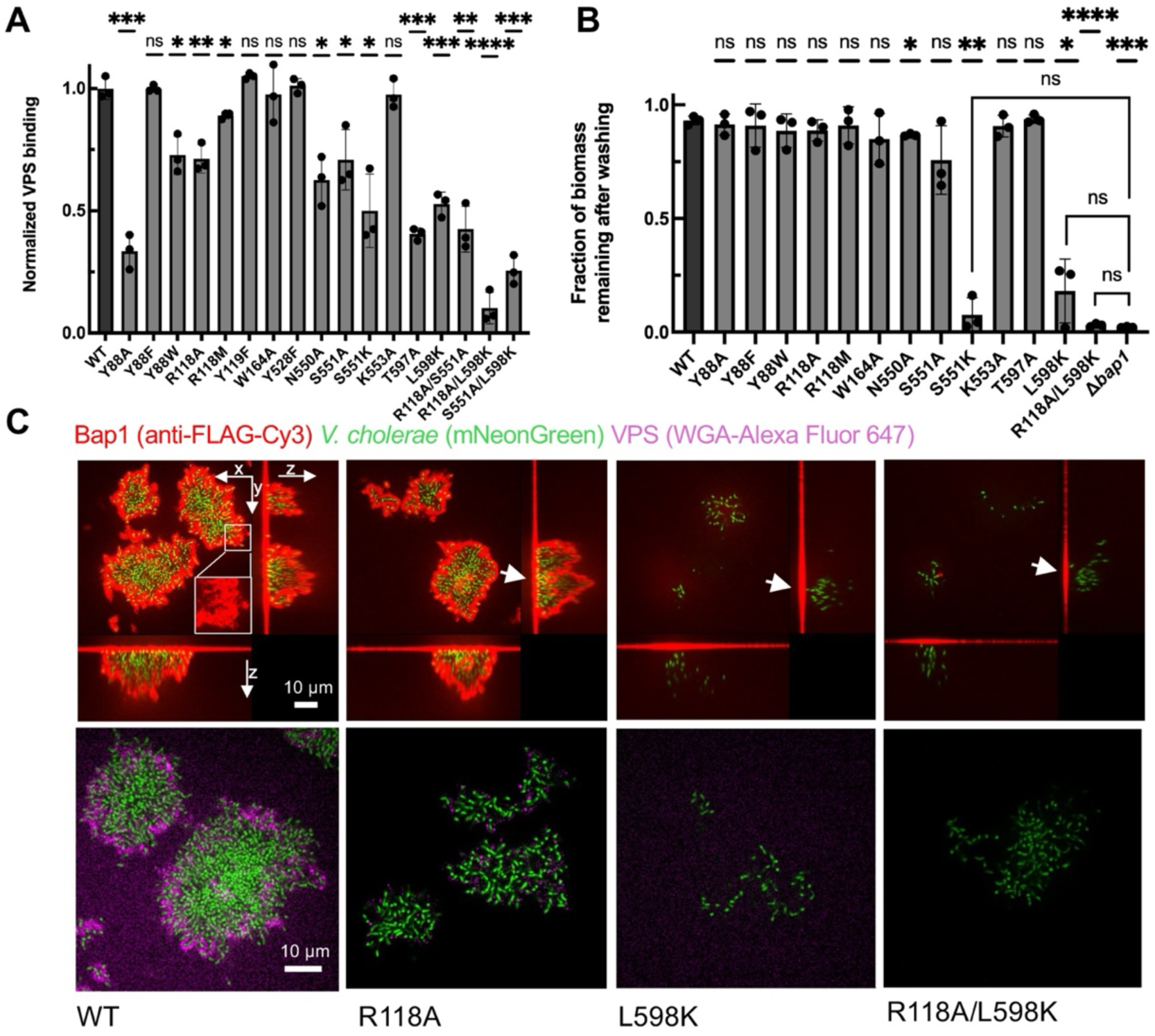
Mutational analysis of VPS-interacting amino acid residues. (A) A native gel-shift analysis showing the disruption of Bap1-VPS interactions of single and double mutations in Bap1. WT data is normalized to 1. (B) Biofilm adhesion assay of *V. cholerae* strains with given mutations introduced into *bap1*. A Δ*rbmC* background is used to avoid the confounding effect of the other adhesin, RbmC. Results are given as the fraction of biofilm mass remaining adhered to the glass substrata following washing. (C) Representative cross-sectional images of biofilms from mutant strains imaged using confocal fluorescence microscopy. *V. cholerae* cells constitutively express mNeonGreen (green); Bap1 proteins tagged with 3×FLAG are stained using an anti-FLAG antibody conjugated to Cy3 (red, *top*), and VPS is stained with wheat germ agglutinin (WGA) conjugated to AlexaFluor 647 (magenta, *bottom*). Arrows indicate secreted Bap1 molecules adhered to the surface, regardless of whether they can bind to VPS and form envelopes surrounding biofilm clusters. Inset shows magnified view of Bap1 staining. Full data for all mutants assayed in (B) are shown in Supplemental Figs. S10-S11. In (A) and (B), statistical significance was determined using an unpaired, two-tailed *t*-test with Welch’s correction compared to WT (unless otherwise indicated by brackets). ns = not significant, *p<0.05, **p<0.01, ***p<0.001, ****p<0.0001.

We made several types of mutations targeting residues that hydrogen-bond directly to VPS, residues that make presumably hydrophobic interactions with VPS, and residues that coordinate the Mg^2+^ ion site interacting with the VPS L-gulose N-acetyl group. In addition to removing putative interactions by mutating residues to alanine, we also made a few mutants to larger side chains, attempting to sterically dislodge VPS from the binding pocket. Starting in numerical order, Y88 forms hydrogen bonds with the L-gulose moiety as well as presumed van der Waals interactions. Mutation of this residue to alanine had a fairly significant effect, reducing the amount of bound Bap1 to approximately 30% (Fig. 4A). Mutation to a more conservative phenylalanine substitution did not significantly reduce binding and mutation to the bulkier tryptophan reduced binding to a more modest 70%. Mutation of R118, which makes several polar interactions with the β-Glc and α-Gal moieties of VPS, to alanine or methionine had minor or insignificant effects. Mutation of Y119 to phenylalanine, which removed a single hydrogen bond with the α-Glc sugar, was also ineffective. Similarly, mutation of Y528 to phenylalanine, and K553 to alanine, did not significantly reduce binding. We were reluctant to mutate the cradling tryptophan residues described in Fig. 1E, reasoning that disrupting these buried hydrophobic residues might have significant effects on the local structure and folding of the β-propeller domain. However, W164, which interacts with the poorly ordered α-Glc group, is less buried than the rest; mutation of this residue to alanine had no significant effect.

We observed more pronounced effects when mutating any of the amino acid sidechains that coordinate the Mg^2+^ ion, including N550A, S551A, and T597A. All three mutations reduced binding to approximately 50% suggesting the importance of this ion binding site in VPS binding. One additional single steric blocking mutation, L598K, was made to disrupt interactions with the L-gulose N-acetyl group resulting in approximately 50% relative binding efficiency. Altogether, these results stress the difficulty of completely disrupting VPS binding with single mutations – presumably, disruption of a single interaction is counteracted by an extensive network of Bap1-VPS interactions encompassing many VPS moieties and Bap1 residues. In an attempt to further disrupt VPS binding, we combined several pairs of single mutants to form double mutants. The combination mutants R118/S551A and S551A/L598K modestly reduced overall binding (∼30-40%), while the combination R118A/L598K demonstrated the most significant overall effect (to ∼10%), also consistent with these being key residues identified in the MD simulations. The dual-pronged approach of dislodging interactions with the L-gulose sugar while releasing interactions with the glucose and galactose groups appeared successful in significantly reducing the interaction between VPS and Bap1 examined by gel-shift analysis.

### Effect of Bap1 mutations on V. cholerae biofilm adhesion and matrix organization

Following *in vitro* characterization of Bap1-VPS interactions, we sought to evaluate the effects of these mutations on the architecture and adhesion of *V. cholerae* biofilms. To remove the compensatory effects of the other β-propeller-containing adhesin RbmC^14^, we used a Δ*rbmC* strain to allow us to solely gauge the contribution of Bap1 mutants to biofilm formation and adhesion. To do this, we used a previously developed biofilm adhesion assay^21^ in which biofilms were grown overnight on glass substrates followed by vigorous washing and quantification of the fraction of remaining biomass by confocal fluorescence microscopy. In this assay, a remaining fraction of 1.0 corresponds to fully functional Bap1 (WT) and 0 corresponds to negative control (Δ*bap1*).

Using this approach we observed a threshold effect, whereby many single mutants that had modest effects in the *in vitro* assay, did not significantly alter biofilm adhesion in this assay (Fig. 4B). This included the mutants Y88F, Y88W, R118A, W164A, K553A, S551A, and N550A. However, the mutants L598K, S551K, and the double mutant R118A/L598K (which previously had been the most disruptive combination mutant) all led to severe adhesion defects statistically indistinguishable from Δ*bap1.* The defective mutants can be rescued through the expression of WT *bap1* from a plasmid (Fig. S8) and we confirmed that the defective phenotype of these *V. cholerae* mutants is not due to defects in secretion or improper folding (Fig. S9). These results confirm our earlier observation that Bap1-VPS interactions are necessary for proper biofilm adhesion to surfaces and, qualitatively agree with the *in vitro* results. Furthermore, they suggest that binding to VPS must be substantially reduced in order to observe defects to biofilm adhesion, possibly arising from the avidity effect that occurs when the entire biofilm attaches to external surfaces.

We reason that some of the milder effects of the mutants may manifest in the mislocalization of the Bap1 molecule or VPS organization in biofilm. Previously, it has been established that Bap1, RbmC, and VPS form envelope structures surrounding *V. cholerae* biofilm clusters, while Bap1 additionally adsorbs spontaneously and strongly to the glass substrate via the 57 amino acid loop^20,21^. Therefore, to further characterize the biofilm morphology of strains expressing mutant Bap1 proteins and to study their localization, we tagged all Bap1 mutants with a 3xFLAG epitope tag and stained secreted Bap1 molecules *in situ* with a fluorescently-labeled anti-FLAG antibody. In the same samples, VPS was additionally visualized using fluorescently-labeled wheat germ agglutinin (WGA), which binds to the minor (20%) GlcNAc component found in VPS. We evaluated all mutants displayed in Fig. 4B, and saw one of three phenotypes (Fig. 4C and Figs. S10-S11). First, we saw mutants with biofilms resembling WT, with Bap1 staining around the periphery of the biofilm and colocalizing with VPS. These mutants were some of those with negligible defects in the gel-shift and adhesion assays including Y88F/W/A, R118M, and W164A (Figs. S10-S11). A second category displayed an intermediate phenotype, with reduced staining of Bap1 and VPS including R118A, N550A, K553A, and T597A. Except for K553A, these mutants showed some reduction in Bap1 binding in the gel-shift assay while all adhered as well as WT in the biofilm adhesion assay.

A third category had the most severe phenotype, with very little Bap1 staining observed in the biofilm, no observable VPS staining by WGA, and a loose cluster of cells attached to the surface. This group included the single mutants S551A, S551K, L598K and the double mutant R118A/L598K, which in general, also displayed defects in Bap1 binding in the gel-shift assay and some of the most severe defects in the adhesion assay. This last group of mutants exhibited substantial substratum-associated Bap1 signals even though little was observed within the biofilm away from the substratum (Fig. 4C and Fig. S10). These results suggest that these mutant Bap1 molecules are still properly secreted and adsorbed to the glass substrate due to their intact 57 amino acid loop, but are unable to engage substantially with the VPS in the biofilm itself. The cells do not completely fall apart likely due to the presence of the cell-cell crosslinking RbmA protein and remaining VPS, but the VPS is in a form not stainable by WGA similar to the negative control (as seen previously in the Δ*bap1*Δ*rbmC* mutant^35^). Altogether, we see a correspondence between mutants judged by *in vitro* binding of VPS by Bap1, biofilm adhesion, and matrix visualization suggesting that the defects seen in the more severe mutations are likely due to disrupted interactions between VPS and Bap1.

### VPS aggregation and effects on biofilm mechanics by Bap1

In addition to mediating adhesion of biofilms to surfaces via interactions with VPS, we have previously observed by native gel-shifts and negative-staining electron microscopy that Bap1 causes the formation of amorphous aggregates of purified VPS when added *in vitro*^21,27^. The presence of only a single VPS binding site in Bap1 as well as no observed Bap1-Bap1 interactions (in crystal packing and in solution) suggests that this is unlikely to be caused by multivalent crosslinking of VPS polymers, but by a yet-to-be-determined mechanism. To further study this aggregation behavior, we used dynamic light scattering (DLS) to characterize the size distribution of VPS polymer during the aggregation process and used the Z-average particle size as a measure of VPS aggregation. In DLS, purified polymeric VPS gives a Z-average of around ∼100 nm (Fig. 5A). Titration of Bap1 into polymeric VPS leads to a shift of the peak to a larger size, near 1,000 nm (Fig. 5A).

**Figure 5.**
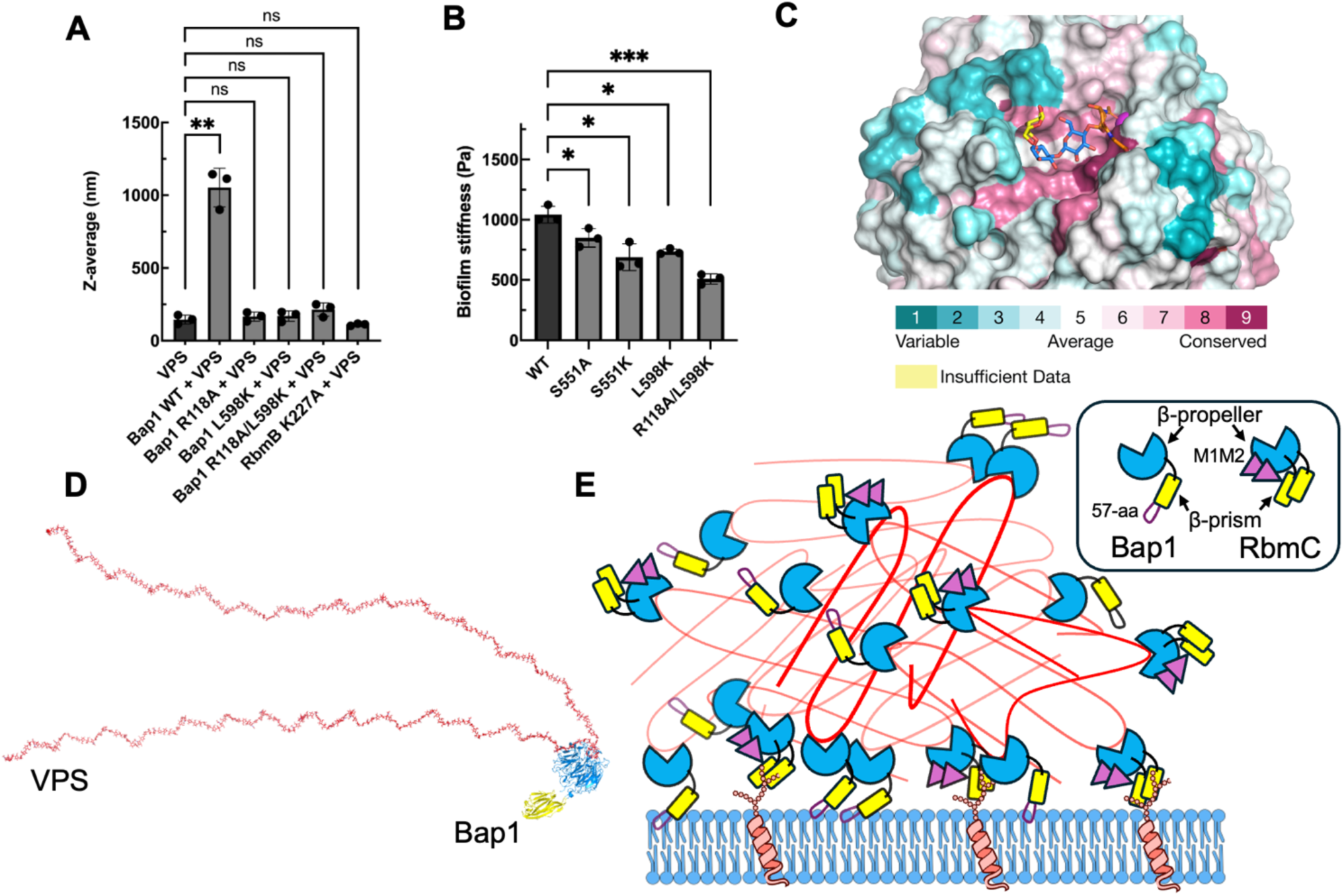
Bap1 mutants are deficient in aggregating VPS. (A) Dynamic light scattering results showing Z-average hydrodynamic sizes (nm) for WT, R118A, L598K, and R118A/L598K show loss of formation of larger aggregates in mutants. (B) Biofilm stiffness, quantified by the plateau modulus G’p, for different mutant biofilms. Statistical significance was determined using an unpaired, two-tailed t-test with Welch’s correction. ns = not significant, *p<0.05, **p<0.01, ***p<0.001. (C) Bap1 surface representation showing sequence conservation of amino acid positions around the VPS binding site in Bap1 and RbmC. Additional details about specific amino acids are shown in Figs. S13 and S14. (D) An 80-mer model highlights the profound impact of the formation of a hairpin kink on the VPS chain. (E) Schematic for VPS compaction caused by bending induced by Bap1 and RbmC. Bap1 adheres to membranes using the 57-aa loop and RbmC uses a combination of N-glycan binding by the β-prism domain and O-glycan-decorated mucin proteins via tandem βγ-crystallin domains (M1M2). Illustration utilizes NIAID NIH BioArt Source (https://bioart.niaid.nih.gov/bioart/468 and https://bioart.niaid.nih.gov/bioart/300).

To determine to what extent VPS-Bap1 interactions observed in the crystal structure are responsible for this aggregation behavior, we used the same DLS-based aggregation assay to assess the effects of the R118A, L598K, and R118A/L598K Bap1 double mutants. None of these mutations showed significant aggregation of VPS in the same concentration range as with the WT Bap1 protein (Fig. 5A, Fig. S12). This suggests that in addition to leading to a defect in substratum adhesion, the Bap1 mutants are also less able to lead to the aggregation of VPS into larger complexes. As a control, we tested the aggregation behavior of a catalytically inactive RbmB^K227A^ mutant that binds VPS and saw that it was unable to aggregate VPS (Fig. 5A).

To test whether the changes in VPS aggregation observed in the light scattering experiment relate to quantitative changes in the physical properties of biofilms, we used shear rheology to measure the stiffness, quantified by the plateau modulus (*G*′_p_ in Pa), of various *V. cholerae* biofilms^36^. All single mutants tested that have a defect in VPS-binding, including S551A, S551K, and L598K mutants, exhibit a significant difference in stiffness as compared to WT biofilms (Fig. 5B). The R118A/L598K double mutant, which displayed more severe losses in binding and adhesion, lead to significantly weaker biofilms, with a stiffness almost half of that of the WT biofilm. These results suggest a direct link between VPS binding, VPS aggregation, biofilm adhesion, and biofilm mechanics mediated by interactions between VPS and Bap1 (and presumably similar for RbmC as well).

### Bioinformatics analysis of VPS binding site in Bap1 and RbmC

Bap1 and RbmC β-propeller domains both bind VPS as judged by native gel-shift assays and share approximately 70% sequence identity between them^21^. Do they both bind to VPS in the same way? What about the β-propeller domains across different *Vibrio* species with homologous biofilm systems^37^? To address these questions, we conducted a bioinformatics analysis to align 107 representative sequences derived by clustering 4,069 Bap1 and RbmC homologs across the *Vibrio* genus (see Materials and Methods). We found a concentration in conserved amino acids around the binding pocket as compared to residues within the rest of the β-propeller domain (Fig. 5C). In fact, many of the amino acid residues involved in Bap1-VPS interactions as identified in our crystal structure were highly conserved within the aligned sequences, both in Bap1 and RbmC (Fig. S13 and S14) suggesting that the VPS binding site is likely conserved across the biofilm-forming species within the *Vibrio* genus. While the possibility of fine-tuning of the binding pocket of the β-propeller in different species is possible, including due to differences in VPS modifications across species, overall we expect the mechanism of VPS binding, aggregation, and adhesion to be conserved throughout these biofilm-forming *Vibrio* species including animal pathogens such as *V. coralliilyticus* and *V. anguillarum*.

## Discussion

Whether to survive fluid flow in the ocean or peristalsis in the gut, biofilms must adhere to surfaces via molecular interactions to diverse substrates and retain sufficient mechanical rigidity. *V. cholerae* does this in a unique way by secreting two adhesins with modular domains directed towards different surface characteristics: RbmC against mucins and *N*-linked glycans and Bap1 against abiotic surfaces and membranes^21^. They also jointly contribute to the mechanical strength of *V. cholerae* biofilms^36,38^. Shared between both adhesins is a homologous β-propeller domain that binds to VPS, an essential and abundant component of the biofilm matrix. Our crystal structure illustrates the molecular details of this interaction to high resolution, illustrating how the β-propeller utilizes a combination of CH-π interactions, hydrogen bonds, hydrophobic interactions, and interactions with a bound ion to bind a single VPS tetrasaccharide. Moreover, we uncovered an unusual bent conformation of VPS when binding takes place, indicating the essential role of conformational shaping of the exopolysaccharide when functioning as the key structural component in the biofilm matrix. Our findings shed light on how a key matrix protein recognizes a cognate exopolysaccharide and how the binding contributes to biofilm integrity and function.

Our structure provides additional context to the unusual chemical modifications found on the L-gulose monosaccharide unit unique to VPS. In the polysaccharide lyase RbmB, the amide-linked glycine moieties play a primary role by forming salt-bridges with arginine sidechains found within the VPS binding groove but no interactions with the N-acetyl groups^13^. In Bap1, the N-acetyl group is tightly enveloped through interactions between the carbonyl oxygen and a bound Mg^2+^ ion, as well as hydrophobic interactions between the N-acetyl methyl group and several hydrophobic and aromatic amino acids. While forming additional hydrogen bond interactions, the amide-linked glycine appears to play a reduced role in Bap1 binding, suggesting that different VPS modifications are uniquely targeted by different VPS binding proteins. These modifications constitute a mechanism for how *V. cholerae* utilizes proteins with drastically different folds to recognize the same polymer. In this regard, it will be interesting to understand how the unrelated yet major matrix protein RbmA, recognizes and binds to VPS.

Through extensive mutagenesis of amino acids that contact VPS in the crystal structure, we confirmed the importance of these interactions in Bap1-VPS binding and in Bap1-mediated adhesion of living biofilms to abiotic substrates. While not completely ruling out the possibility for additional VPS binding sites, disruption of this Bap1-VPS binding interface leads to a series of defects including biofilm surface adhesion, matrix organization, and biofilm mechanics suggests that this interface is a critical component of the adhesion mechanism. We found the Bap1 binding site challenging to disrupt with single point mutations, likely due to the adhesin’s high affinity for VPS. Even when we observed some loss of VPS binding *in vitro*, these mutants had weak effects in biofilm adhesion assays. We did, however, identify several single (S551A, S551K, and L598K) and a combination double mutant (R118A/L598K) that displayed severe defects in functional assays.

Unexpectedly, our structure also indicated that the VPS tetramer is bent severely within the Bap1 binding site, with the central D-Glcβ(1-4)-D-Glcα bond occupying a high-energy state. Note that this glycosidic bond is the same as the main glycosidic bond in cellulose, whose lowest energy state leads to a linear conformation-the same low-energy state observed in the RbmB-VPS crystal structure and in the free VPS polymer by MD simulations^13^. Whether this results from Bap1-induced bending of the VPS polymer as Bap1 binds, or conformational capture of already bent VPS polymers in solution is not certain. Either way, this bending is quite unusual for known carbohydrate polymers binding to proteins, which typically evolve to recognize the lowest energy state of the polymer as seen with free polymer^39,40^. The overall effect of this binding is illustrated in an 80-mer VPS model, indicating that Bap1 creates a hairpin structure in VPS, reversing the direction of the VPS chain in solution (Fig. 5D).

In addition to acting as “double-sided tape” in adhering biofilms to surfaces^21^, our DLS experiments suggest an independent mechanism for Bap1 in aggregating VPS into large amorphous assemblies through the β-propeller (Fig. 5E). This aggregation is tightly related to proper VPS organization in biofilm clusters and the mechanical strength of the biofilm. The presence of a single VPS binding site in Bap1 argues against the action of Bap1 as a multivalent crosslinker, requiring a different mechanistic understanding for this activity. One possibility is that monovalent Bap1-VPS interactions induce weak Bap1-Bap1 interactions leading to condensation of Bap1-coated polymers into larger aggregates, however, we have never observed Bap1 in any other oligomeric state. Another possibility is that Bap1 binding of VPS alters the solubility of VPS, leading to precipitation into larger aggregates. For example, binding of Bap1 could mask VPS negative charges or by increasing VPS-VPS interactions through bending of the polymer into a more “sticky” state. The observation that a catalytically inactive RbmB^K227A^ mutants that binds VPS, but that only bends VPS by 30°^13^, leads to much less aggregation favors the latter possibility. However, additional studies delineating the complex interactions between these proteins and VPS are necessary to reach a unified mechanism for VPS aggregation and function. Modeling of longer chains of VPS bound to Bap1 suggests that Bap1 creates the possibility of hairpin-like structures in VPS, a motif not seen in glycans except by design^41^. How this large distortion of the VPS structure leads to aggregation behavior will continue to be a focus of ongoing studies.

Together, our structure provides a high-resolution view into the molecular interaction between a *V. cholerae* matrix protein and the primary biofilm matrix exopolysaccharide. This interaction underlies many of the *V. cholerae* biofilm properties, including surface adhesion, cell ordering, proper localization and maintenance of exopolysaccharides, and biofilm rigidity. Understanding how biofilms are assembled is not only essential for targeting bacterial pathogens of medical and agricultural interest, but also in designing new biomaterials that may serve as underwater glues with medical and industrial applications – our study provides a template for the study of such interactions in many biological systems.

## Methods and Materials

### Recombinant Bap1 expression and purification

All Bap1 constructs were cloned, expressed and purified in *E. coli* as previously described^22^, with the following exception: site-directed mutagenesis was used to remove the thrombin cleavage site on the linker separating the His_6_-GFP_UV_ tag from the insert in the N-terminus of the vector pNGFP-BC to improve protein yield during expression and purification^21^. All *E. coli*-purified proteins and mutants were derived from Bap1_Δ57_ (referred to throughout as Bap1), which is missing the 57-amino acid adhesion loop to retain solubility^22^. Trypsin was used to cleave the His_6_-GFP tag from the Bap1 proteins as follows: a 1:500 protein to trypsin ratio and 1 mL digestion volume was incubated at room temperature for 2 hours in the presence of 10 mM CaCl_2_. The trypsin digestion was stopped by 1 mM AEBSF and followed by centrifugation at 21,000 x g for 5 minutes. Supernatants were applied to a Q-ion exchange chromatography column as previously described^22^. Site-directed mutagenesis was used to produce all single and double Bap1 mutants in this study using overlapping PCR primers (Table S9). To further enhance Bap1 protein expression, a double colony selection protocol^42^ was followed for WT Bap1 – this formed the template to produce all Bap1 mutants via site-directed mutagenesis.

### Vibrio cholerae strains and growth

All *V. cholerae* strains used in this study were derivatives of the WT *V. cholerae* O1 biovar El Tor strain C6706str2 and listed in Table S10. The rugose strain background harbors a missense mutation in the *vpvC* gene (*vpvC*^W240R^) that elevates intracellular cyclic diguanylate levels^43^. The rugose strains form robust biofilms and thus allow us to focus on the biochemical mechanisms governing biofilm adhesion rather than mechanisms involving gene regulation. *rbmC* was deleted from the strain to focus on the effect of mutations in *bap1* on VPS binding. Additional mutations were genetically engineered into this *V. cholerae* strain using the natural transformation (MuGENT) method or through conjugation^44^.

All strains were grown overnight in lysogenic broth (LB) at 37 °C with shaking. 1× M9 media were filter sterilized and supplemented with 2 mM MgSO_4_ and 100 µM CaCl_2_ (abbreviated as M9 medium below). Biofilm growth was generally performed in M9 medium supplemented with 0.5% glucose. For complementation experiments, 100 µg/mL kanamycin was used to maintain the plasmid.

### Vibrio cholerae strain construction

Linear PCR products were constructed using splicing-by-overlap extension (SOE) PCR as previously described and used as transforming DNA (tDNA) in chitin-dependent transformation reactions^44^. Briefly, SOE PCR was performed by amplifying an upstream region of homology and a downstream region of homology. The desired mutations were incorporated into the primers used in amplification. All primers used to construct and detect mutant alleles are listed in Table S9. For chitin-dependent transformation, individual *V. cholerae* colonies were grown in LB media at 30 °C for 6 h to an OD_600_ = 0.8–1.0. Cells were washed with Instant Ocean (IO) solution and then incubated with chitin particles suspended in IO for 8–16 h at 30 °C before the tDNA was added. The cultures were then incubated at 30 °C for an additional 8–16 h. LB was added to the cultures and incubated at 37 °C for 2 h before plating on LB agar with the appropriate antibiotic. The desired mutants were selected by the emergence of new phenotype or colony PCR screening and confirmed by sequencing and complementation.

### Purification of VPS fragments

VPS was extracted and purified from *V. cholerae* biofilms as previously described with the only modification that LB were used as a growth media to increase the yield^21^. To produce VPS fragments for crystallization, purified VPS polymers were digested by recombinant RbmB^27^ at 1:10 weight ratio for ∼72 hours at 30 °C. Digested VPS was cleaned up to remove RbmB by boiling for 10 min before purifying using solid phase extraction^27^. SPE eluates were lyophilized via speed-vac for ∼2 hours and resuspended in <100 μL volumes of SEC buffer to produce final digested VPS concentrations of 1-10 mM as measured by borate-ethanolamine fluorescence using a standard curve of galactose^45,46^. This concentrated digested VPS was used as a direct substrate for Bap1 crystallography.

### Crystallization of VPS–Bap1

Bap1 crystals were reproduced and flash-frozen as previously described^22^, with some modifications. Briefly, Bap1 was concentrated to 5 mg/mL, VPS was added to a final concentration of approximately 0.7 mM digested VPS, and crystallized by the hanging drop method in 1:1 drops using a buffer containing 12% PEG 3,350, 100 mM sodium citrate, pH 5.4. For cryoprotection before data collection, crystals were soaked in mother liquor containing 20% glycerol and 0.7 mM digested VPS to prevent substrate ligand back-soaking and flash-frozen in liquid nitrogen. Crystals were of a similar morphology as the native crystals reported previously^22^.

### X-ray structure determination and refinement of VPS–Bap1

VPS–Bap1 X-ray data were collected remotely at the Brookhaven National Laboratory NSLS-II beamline 17-ID-2 (FMX) and processed using the autoPROC^47^ pipeline (including the programs XDS^48^, POINTLESS^49^, AIMLESS^50^, and CCP4^51^). Data were elliptically truncated using STARANISO^52^. The crystals were isomorphous with the previously solved apo Bap1 structure^22^ allowing the previous coordinates (PDB:6MLT) to be used as a starting model for refinement. Multiple rounds of positional and B-factor refinement with TLS were carried out using phenix.refine^53^ with anisotropic B-factors used for the metals and VPS. The CheckMyMetal website (https://cmm.minorlab.org/)^54^ was used to identify and validate bound calcium and magnesium ions. Models were rebuilt using COOT^55^ using unfilled maps (due to the elliptical truncation) and final figures generated using PyMOL^34^. Model restraints for the modified L-gulose carbohydrate ligand were generated using eLBOW^56^ in the Phenix structure determination package^57^. For figures, Polder^30^ omit maps were generated to display the VPS ligand. To remove possible phase bias before generating these maps, the carbohydrate ligand was removed, all B-factors were set to a value of 20 Å^2^, and all atomic coordinates displaced 0.1 Å in random directions using PDBSET in CCP4^51^. The resulting coordinates were then re-refined with five cycles of positional and isotropic B-factor refinement without the carbohydrate ligands before calculating maps in PHENIX^57^.

### Native PAGE shift assays

Tris-glycine native gels containing 4% and 10% acrylamide stacking and resolving layers, respectively, were run in ice at 80 V for 4-6 hours until desired separation was achieved^21^. Protein preps of WT or mutant Bap1-GFP_UV_ fusion were first purified via a Q-sepharose ion exchange column prior to Native PAGE. To quantify relative aggregation activity of polymeric VPS, 10 µg of polymeric VPS was mixed with 5 µg protein 5 minutes prior to loading the Native PAGE. VPS-induced gel shift of the fluorescence protein band corresponding to Bap1-GFP_UV_ protein was imaged using a Typhoon FLA 9000 flatbed imager (GE/Cytiva) using the blue laser (473 nm).

Pixel quantification of Bap1-GFP_UV_ bands was performed on protein-only gel bands using ImageQuant TL v. 7.0. The volume of protein loaded for every gel lane was determined by absorbance at 280 nm while taking account for % purity estimated by SDS PAGE. Bap1 WT was loaded as a positive control for every native gel, which also featured 1-2 mutants in every native gel. GFP fluorescence, and thus, binding activity was normalized to WT ±VPS. Consequently, every apo (-VPS) protein exhibited ±30% pixel intensity across all proteins within a single native gel. The change in pixel density for mutant Bap1 proteins was normalized to the activity of a WT protein sample measured on the same gel as each mutant protein. Biological replicates of three were performed for each mutant to produce a pseudo-quantitative VPS-binding activity relative to WT.

### Dynamic light scattering titration assays to quantify Bap1-VPS aggregation

DLS experiments were conducted using a Horiba SZ-100-Z2 instrument (Horiba, CA). Following trypsin cleavage of the N-terminal His_6_-GFP_UV_ tag and Q-ion exchange chromatography, concentrated Bap1 WT and mutant protein stocks were centrifuged at 21,000 x g for 20 minutes to pellet large particles. Supernatants were diluted to 0.3-0.5 mg/mL and filtered through PVDF 0.22 μm syringe filter. Proteins were incubated with 1:1 mass VPS or buffer volume for 30 minutes prior to DLS. Horiba software (FluorEssence v. 3.9.0.1) was used to quantify relative changes to hydrodynamic radii size (Z-average) via fitting of the autocorrelation function. Z-average values for all samples were produced by averaging 3 measurements of 3 technical replicates (9 total measurements). DLS settings were as follows: auto scattering mode (90° or side scatter), 60 s scan time, monodisperse fitting mode of normal peak width.

### Biofilm adhesion assay

Overnight cultures of the indicated strains constitutively expressing mNeonGreen were grown from individual colonies at 37°C with shaking in 1.5 mL LB. 50 µL from each culture was used to inoculate 1.5 mL of M9 medium supplemented with 0.5% glucose and grown at 30°C with shaking until the OD_600_ was between 0.1 and 0.3. The cultures were then diluted to an OD_600_ ≅ 0.001. 100 μL of the regrown culture was aliquoted into the wells of a 96-well plate with a glass bottom (MatTek P96G-1.5-5-F) and incubated at 30°C for 1 h. The wells were then washed twice with M9 medium and replaced with M9 medium with 0.5% glucose. In the complementation experiment, the growth medium additionally contains 0.2% arabinose for P_BAD_-*bap1*. The lid was secured with a layer of parafilm and the 96-well plate was subsequently incubated at 30°C for 16–24 h. Thus-prepared samples were imaged with a spinning disk confocal microscope (Nikon Ti2-E connected to Yokogawa W1) using a 60× water objective (numerical aperture = 1.20) and a 488 nm laser excitation. For each sample, several locations with 3 × 3 tiles where imaged and captured with an sCMOS camera (Photometrics Prime BSI or Hamamatsu Fusion BT). The *x*-*y* pixel size was 0.22 μm and the *z*-step size was 3 μm. The wells were then washed twice with M9 medium and re-imaged at the same locations. All images presented in this study are raw data rendered using the Nikon Elements software.

Image analysis was performed with built-in functions of the Nikon Elements software by thresholding each image layer-by-layer and measuring the total binarized area above the threshold in each layer. The binary area for each sample *z*-slice was then summed to give the total biovolume, and the ratio of the total biovolume after versus before the washing step was calculated.

### In situ biofilm immunostaining

Overnight cultures of the indicated strains with WT or mutated *bap1* tagged with 3×FLAG at the C-terminus and constitutively expressing mNeonGreen were grown following the same procedure as described above. The initial incubation time was adjusted to 30 min. The wells were washed twice with M9 medium; subsequently, 100 µL of M9 medium with 0.5% glucose and 0.5 mg/mL BSA (Sigma-Aldrich A9647) was added to the well. The growth media additionally included 4 µg/mL anti-FLAG antibody conjugated to Cy3 (Sigma-Aldrich A9594) and 4 µg/mL WGA conjugated to AlexaFluor647 (W32466 ThermoFisher). The lid was secured with a layer of parafilm and incubated at 30 °C for 16–24 h. Thus-prepared samples were carefully mounted onto a spinning disk confocal microscope (Nikon Ti2-E connected to Yokogawa W1) using a 100× oil immersion objective (numerical aperture = 1.35) and a 488 nm laser excitation to observe the cells and a 561 nm laser excitation to observe protein localization, with the corresponding filters. The images were captured with an sCMOS camera (Hamamatsu Fusion BT) at a *z*-step size of 0.5 µm, going from 2 µm below the glass to ∼ 30 µm above the glass to cover the entire biofilm in the imaging volume. Subsequently, the medium in each well was gently removed and replaced with fresh M9 medium, and the samples were imaged again on the spinning disk with a 488 nm laser excitation to observe the cells and a 640 nm laser to observe the WGA staining of VPS. We observed that biofilms from some mutants with adhesion defects will move during the medium replacement step.

### Western blots and secretion assay

*V. cholerae* strains encoding the indicated constructs with a C-terminal 3×FLAG tag were grown in culture tubes containing 3 mL LB overnight at 30 °C. The next day, cultures were vortexed to break up pellicles and cell clusters and the OD_600_ was measured. Dilution of each cell suspension to the lowest optical density measured was conducted in sterile 1.5 mL microcentrifuge tubes and spun at 18,000 × g for 1 min so that each tube would have 200 μL total. ∼200 μL of the cell supernatant was transferred to a fresh 1.5 mL microcentrifuge tube. The cell pellets were lysed for 20 min using a lysis solution (1× Bugbuster solution, lysozyme (0.05-0.1 mg/mL), and benzonase (≥ 250 units/mL)). 30 μL of each cell suspension was combined with 10 μL of 4× SDS PAGE sample buffer (40% Glycerol, 240 mM Tris pH 6.8, 8% SDS, 0.04% Bromophenol Blue, 5% β-mercaptoethanol) and boiled for 10 min at 95 °C. Samples were run on a 4– 15% Mini-PROTEAN TGX gel in 1× SDS PAGE running buffer (25 mM Tris, 192 mM Glycine, 1% SDS, pH 8.3) at 150 V for 60 min. The proteins were transferred to a PVDF membrane in 1× Transfer buffer (25 mM Tris, 192 mM Glycine, 10% methanol, pH 8.3) at 100 V for 1 h. The membranes were incubated in 5% milk in TBST for 1 h at room temperature. The membranes were washed 3 × 10 min in 1× TBST. The membranes were blotted using α-DYKDDDDK (BioLegend 637311) at 0.1 μg/mL in TBST with 3% BSA for 1 h at room temperature and washed 3 × 10 min with 1× TBST. Blots were developed by incubation with Super Signal PLUS Pico West Chemiluminescent Substrate for 5 min and pictures taken using the BioRad Chemidoc-MP. The membranes were washed 3 × 10 min in 1× TBST and probed with anti-RpoB-HRP antibody solution. Blots were again developed by incubation with Super Signal PLUS Pico West Chemiluminescent Substrate for 5 min and pictures taken using the BioRad Chemidoc-MP.

### Rheology

The rheological measurement follows a previously described procedure^36^. *V. cholerae* strains (in a ΔpomA background) were streaked on LB plates containing 1.5% agar and grown at 37°C overnight. Individual colonies were inoculated into 3 mL of LB liquid medium containing glass beads, and the cultures were grown with shaking at 37°C to mid-exponential phase (5-6 h). Subsequently, the cells in the cultures were mixed by vortex, OD600 was measured, and the cultures were back diluted to an OD_600_ of 0.5. 1 μL of this inoculum was spotted onto pre-warmed agar plates (100 mm) solidified with 0.6% LB agar. Each plate contains multiple colony biofilms, and multiple plates were prepared for each rheological measurement. All rheological measurements were performed with a stress-controlled Anton Paar Physica MCR WESP502 rheometer. The colony biofilms were collected with a pipette tip or a razor blade and transferred onto the lower plate of the rheometer. After sandwiching the biofilms between the upper and lower plates with a gap size of 0.5 mm, silicone oil (5 cSt at 25°C, Sigma Aldrich) was applied to surround the biofilm to avoid evaporation. Sandblasted surfaces were used for both the upper and lower plates to avoid slippage at the boundary. Oscillatory shear tests were performed. During amplitude sweeps, a strain range of 0.01 – 2000% was scanned at a fixed frequency of 6.28 rad s^−1^ at 37°C. Segmented linear fittings were applied to *G*’ curves on a log-log scale to find the plateau modulus *G*’_p_^58^ (stiffness).

### Modeling and molecular dynamics of the Bap1-VPS complex

MD simulations were performed using the Bap1–VPS co-crystal structure as the starting model to investigate the molecular interactions governing Bap1–VPS binding across multiple complex variants, including complexes containing tetrameric and dodecameric VPS fragments. The dodecameric substrate model was constructed by extending four residues on both ends using the GLYCAM-Web^59^ (glycam.org) models as a template. Using the Amber software tool tLEaP^60^, counter ions were added to neutralize the system and the 3D structures were placed in a truncated octahedron box of TIP5P water with an 8 Å water buffer. Non-bonded cutoffs of 8 Å for van der Waals and electrostatics were employed. All MD simulations were performed with the Amber24 software using the GLYCAM-06j force field^61^ for carbohydrates and the ff19SB force field^62^ for the protein. Energy minimization, heating, equilibration and production MD were performed according to recommended protocols^63^. Production MD data were collected for a total trajectory time of 0.5 µs for each system at constant pressure (1 atm) and room temperature (298 K). Snapshots were recorded every 100 ps during MD simulation for downstream analysis using the AMBER CPPTRAJ program^64^. The carbohydrate intrinsic (CHI) energies associated with glycosidic 1-4 linkages were utilized to analyze the dihedral angle distributions of the glycosidic linkages^40^. The MM-GBSA method^65^ was used to quantify the per-residue contributions to the overall interaction energy, employing the GBHCT parameterization (IGB=2) with the mbondi2 Born radii set.

### Bioinformatics analysis of Bap1/RbmC across Vibrio species

All genomes assigned to the genus *Vibrio* were retrieved from the Genome Taxonomy Database (GTDB, release 214^66^), and protein-coding sequences were annotated using Prokka^67^ (version 1.14.6). Of the 1,983 genomes classified as *Vibrio*, only those with estimated completeness ≥90% and contamination ≤5%, as determined by CheckM^68^, were retained for further analysis. Homologs of *rbmC* and *bap1* were identified using the following quality filters: ≥80% amino acid identity to a *bap1* query sequence, sequence length between 650 and 700 amino acids, and bit scores greater than 900. To reduce redundancy, protein sequences were clustered with CD-HIT^69^ (version 4.8.1) at 98% sequence identity, yielding 107 representative sequences from an initial dataset of 4,069. Representative sequences were aligned with MAFFT^70^ (version 7.526) using the options “--maxiterate 1000--localpair.” Sequence conservation was assessed with Consurf^71^ (https://consurf.tau.ac.il/consurf_index.php), which calculates conservation scores based on sequence variability across the alignment. Residues within 8 Å of the glycan ligand were designated as potential binding sites, resulting in the identification of 55 residues, which were subsequently plotted for visualization.

### Statistics and reproducibility

Error bars correspond to standard deviations from measurements taken from distinct samples. Standard *t*-tests were used to compare treatment groups and are indicated in each figure legend. Tests were always two-tailed, unpaired, and used Welch’s correction, as demanded by the details of the experimental design. All statistical analyses were performed using GraphPad Prism v. 10.5.0 software. Microscopy images were shown from representative results from at least three independent experiments.

## Acknowledgements

We thank Drs. Lei Li and Fitnat Yildiz for helpful discussions. We thank Drs. James Murphy, Titus Boggon, and Kathryn Ferguson for assistance in obtaining beamtime at BNL. We thank Dr. Craig Crews, Dr. Mackenzie Krone, and Dr. Chunxiang Wu for sharing equipment and for assistance in data collection. Research reported in this publication was supported by the National Institute of General Medical Sciences of the National Institutes of Health (https://www.nigms.nih.gov/) under award number 1DP2GM146253 to J.Y. and R15GM152959 to R.O. J.Y. acknowledges support from the National Science Foundation (#2205006), Army Research Office (W911NF-25-1-0084), Alfred P. Sloan Foundation (FG-2023-20857), and the Burroughs Wellcome Foundation (#1022835). Y.Y and X.J. are supported by the Division of Intramural Research of the NIH, National Library of Medicine. This work utilized the computational resources of the NIH HPC Biowulf cluster (http://hpc.nih.gov). This research used beamline 17-ID-2 (FMX) of the National Synchrotron Light Source II, a U.S. Department of Energy (DOE) Office of Science User Facility operated for the DOE Office of Science by Brookhaven National Laboratory under Contract No. DE-SC0012704. The Center for BioMolecular Structure (CBMS) is primarily supported by the National Institutes of Health, National Institute of General Medical Sciences (NIGMS) through a Center Core P30 Grant (P30GM133893), and by the DOE Office of Biological and Environmental Research (KP1607011). Beamtime was obtained through the Yale University Block Allocation Group #317429. This work was supported in part by GlycoMIP, a National Science Foundation Materials Innovation Platform funded through Cooperative Agreement DMR-1933525 (RJW).

## References

1. Flemming, H.-C. et al. Biofilms: an emergent form of bacterial life. Nat. Rev. Microbiol. 14, 563–575 (2016).

2. Jiang, Z., Nero, T., Mukherjee, S., Olson, R. & Yan, J. Searching for the secret of stickiness: how biofilms adhere to surfaces. Front. Microbiol. 12, (2021).

3. Flemming, H.-C. et al. The biofilm matrix: multitasking in a shared space. Nat. Rev. Microbiol. 21, 70–86 (2023).

4. Essentials of Glycobiology. (Cold Spring Harbor Laboratory Press, Cold Spring Harbor (NY), 2015).

5. Fisher, R. A., Gollan, B. & Helaine, S. Persistent bacterial infections and persister cells. Nat. Rev. Microbiol. 15, 453–464 (2017).

6. Niu, H., Gu, J. & Zhang, Y. Bacterial persisters: molecular mechanisms and therapeutic development. Signal Transduct. Target. Ther. 9, 174 (2024).

7. Nelson, E. J., Harris, J. B., Morris, J. G., Calderwood, S. B. & Camilli, A. Cholera transmission: the host, pathogen and bacteriophage dynamic. Nat. Rev. Microbiol. 7, 693–702 (2009).

8. World Health Organization. Cholera, 2024. Wkly. Epidemiol. Rec. 100, 347–363 (2025).

9. Teschler, J. K. et al. Living in the matrix: assembly and control of *Vibrio cholerae* biofilms. Nat. Rev. Microbiol. 13, 255–268 (2015).

10. Fong, J. N. C. & Yildiz, F. H. Biofilm matrix proteins. Microbiol. Spectr. 3, (2015).

11. Teschler, J. K., Nadell, C. D., Drescher, K. & Yildiz, F. H. Mechanisms underlying *Vibrio cholerae* biofilm formation and dispersion. Annu. Rev. Microbiol. 76, 503–532 (2022).

12. Yildiz, F., Fong, J., Sadovskaya, I., Grard, T. & Vinogradov, E. Structural characterization of the extracellular polysaccharide from *Vibrio cholerae* O1 El-Tor. PLoS ONE 9, e86751 (2014).

13. Weerasekera, R. et al. Crystal structure of *Vibrio cholerae* polysaccharide lyase RbmB bound to *Vibrio* polysaccharide (VPS) fragments provides insights into substrate recognition and cleavage. Proc. Natl. Acad. Sci. 123, e2534280123 (2026).

14. Fong, J. C. N. & Yildiz, F. H. The *rbmBCDEF* gene cluster modulates development of rugose colony morphology and biofilm formation in *Vibrio cholerae*. J. Bacteriol. 189, 2319–2330 (2007).

15. Fong, J. C. N., Karplus, K., Schoolnik, G. K. & Yildiz, F. H. Identification and characterization of RbmA, a novel protein required for the development of rugose colony morphology and biofilm structure in *Vibrio cholerae*. J. Bacteriol. 188, 1049– 1059 (2006).

16. Giglio, K. M., Fong, J. C., Yildiz, F. H. & Sondermann, H. Structural basis for biofilm formation via the *Vibrio cholerae* matrix protein RbmA. J. Bacteriol. 195, 3277–3286 (2013).

17. Maestre-Reyna, M., Wu, W.-J. & Wang, A. H.-J. Structural insights into RbmA, a biofilm scaffolding protein of *V. cholerae*. PLoS ONE 8, e82458 (2013).

18. Fong, J. C. et al. Structural dynamics of RbmA governs plasticity of *Vibrio cholerae* biofilms. eLife 6, e1002210 (2017).

19. Absalon, C., Van Dellen, K. & Watnick, P. I. A communal bacterial adhesin anchors biofilm and bystander cells to surfaces. PLoS Pathog. 7, e1002210 (2011).

20. Berk, V. et al. Molecular architecture and assembly principles of *Vibrio cholerae* biofilms. Science 337, 236–239 (2012).

21. Huang, X., et al. *Vibrio cholerae* biofilms use modular adhesins with glycan-targeting and nonspecific surface binding domains for colonization. Nat. Commun. 14, 2104 (2023).

22. Kaus, K. et al. The 1.9 Å crystal structure of the extracellular matrix protein Bap1 from *Vibrio cholerae* provides insights into bacterial biofilm adhesion. J. Biol. Chem. 294, 14499–14511 (2019).

23. Huang, X. et al. Conformations and sequence determinants in the lipid binding of an adhesive peptide derived from Vibrio cholerae biofilms. PLOS Pathog. 22, e1013990 (2026).

24. De, S., Kaus, K., Sinclair, S., Case, B. C. & Olson, R. Structural basis of mammalian glycan targeting by *Vibrio cholerae* cytolysin and biofilm proteins. PLoS Pathog. 14, e1006841 (2018).

25. Jaroentomeechai, T. et al. Microbial binding module employs sophisticated clustered saccharide patches to selectively adhere to mucins. Nat. Commun. 16, 9058 (2025).

26. Barrasso, K. et al. Impact of a human gut microbe on *Vibrio cholerae* host colonization through biofilm enhancement. eLife 11, e73010 (2022).

27. Weerasekera, R., et al. *Vibrio cholerae* RbmB is an α-1,4-polysaccharide lyase with biofilm-disrupting activity against *Vibrio* polysaccharide (VPS). PLOS Pathog. 20, e1012750 (2024).

28. Bridges, A. A., Fei, C. & Bassler, B. L. Identification of signaling pathways, matrix-digestion enzymes, and motility components controlling *Vibrio cholerae* biofilm dispersal. Proc. Natl. Acad. Sci. 117, 32639–32647 (2020).

29. Wardman, J. F., Bains, R. K., Rahfeld, P. & Withers, S. G. Carbohydrate-active enzymes (CAZymes) in the gut microbiome. Nat. Rev. Microbiol. 20, 542–556 (2022).

30. Liebschner, D. et al. Polder maps: improving OMIT maps by excluding bulk solvent. Acta Crystallogr. Sect. Struct. Biol. 73, 148–157 (2017).

31. Hudson, K. L. et al. Carbohydrate–aromatic interactions in proteins. J. Am. Chem. Soc. 137, 15152–15160 (2015).

32. Laskowski, R. A. & Swindells, M. B. LigPlot+: multiple ligand-protein interaction diagrams for drug discovery. J. Chem. Inf. Model. 51, 2778–2786 (2011).

33. Krissinel, E. & Henrick, K. Inference of macromolecular assemblies from crystalline state. J. Mol. Biol. 372, 774–797 (2007).

34. Schrödinger, LLC. The PyMOL molecular graphics system, version 3.1.6.1.

35. Yan, J., Sharo, A. G., Stone, H. A., Wingreen, N. S. & Bassler, B. L. Vibrio cholerae biofilm growth program and architecture revealed by single-cell live imaging. Proc. Natl. Acad. Sci. 113, E5337–E5343 (2016).

36. Yan, J. et al. Bacterial biofilm material properties enable removal and transfer by capillary peeling. Adv. Mater. 30, 1804153 (2018).

37. Yang, Y., Yan, J., Olson, R. & Jiang, X. Comprehensive genomic and evolutionary analysis of biofilm matrix clusters and proteins in the *Vibrio* genus. mSystems 0, e00060–25 (2025).

38. Zhang, Q., et al. Mechanical resilience of biofilms toward environmental perturbations mediated by extracellular matrix. Adv. Funct. Mater. 32, 2110699 (2022).

39. Boraston, A. B., Bolam, D. N., Gilbert, H. J. & Davies, G. J. Carbohydrate-binding modules: fine-tuning polysaccharide recognition. Biochem. J. 382, 769–781 (2004).

40. Nivedha, A. K., Makeneni, S., Foley, B. L., Tessier, M. B. & Woods, R. J. Importance of ligand conformational energies in carbohydrate docking: Sorting the wheat from the chaff. J. Comput. Chem. 35, 526–539 (2014).

41. Fittolani, G. et al. Synthesis of a glycan hairpin. Nat. Chem. 15, 1461–1469 (2023).

42. Sivashanmugam, A. et al. Practical protocols for production of very high yields of recombinant proteins using *Escherichia coli*. Protein Sci. 18, 936–948 (2009).

43. Beyhan, S. & Yildiz, F. H. Smooth to rugose phase variation in *Vibrio cholerae* can be mediated by a single nucleotide change that targets c-di-GMP signalling pathway. Mol. Microbiol. 63, 995–1007 (2007).

44. Dalia, A. B., McDonough, E. & Camilli, A. Multiplex genome editing by natural transformation. Proc. Natl. Acad. Sci. 111, 8937–8942 (2014).

45. Kato, T. & Kinoshita, T. Fluorometric detection and determination of carbohydrates by high-performance liquid chromatography using ethanolamine. Anal. Biochem. 106, 238–243 (1980).

46. O’Neill, R. A., Darvill, A. & Albersheim, P. A fluorescence assay for enzymes that cleave glycosidic linkages to produce reducing sugars. Anal. Biochem. 177, 11–15 (1989).

47. Vonrhein, C. et al. Data processing and analysis with the autoPROC toolbox. Acta Crystallogr. D Biol. Crystallogr. 67, 293–302 (2011).

48. Kabsch, W. XDS. Acta Crystallogr. D Biol. Crystallogr. 66, 125–132 (2010).

49. Evans, P. Scaling and assessment of data quality. Acta Crystallogr. D Biol. Crystallogr. 62, 72–82 (2006).

50. Evans, P. R. & Murshudov, G. N. How good are my data and what is the resolution? Acta Crystallogr. D Biol. Crystallogr. 69, 1204–1214 (2013).

51. Winn, M. D. et al. Overview of the CCP4 suite and current developments. Acta Crystallogr. D Biol. Crystallogr. 67, 235–242 (2011).

52. Tickle, I.J., Flensburg, C., Keller, P., Paciorek, W., Sharff, A., Vonrhein, C., Bricogne, G. STARANISO. Cambridge, United Kingdom: Global Phasing Ltd. Global Phasing Ltd. (2018).

53. Afonine, P. V. et al. Towards automated crystallographic structure refinement with phenix.refine. Acta Crystallogr. D Biol. Crystallogr. 68, 352–367 (2012).

54. Zheng, H. et al. Validation of metal-binding sites in macromolecular structures with the CheckMyMetal web server. Nat. Protoc. 9, 156–170 (2014).

55. Emsley, P., Lohkamp, B., Scott, W. G. & Cowtan, K. Features and development of Coot. Acta Crystallogr. D Biol. Crystallogr. 66, 486–501 (2010).

56. Moriarty, N. W., Grosse-Kunstleve, R. W. & Adams, P. D. Electronic ligand builder and optimization workbench (eLBOW): a tool for ligand coordinate and restraint generation. Acta Crystallogr. D Biol. Crystallogr. 65, 1074–1080 (2009).

57. Adams, P. D. et al. PHENIX: a comprehensive python-based system for macromolecular structure solution. Acta Crystallogr. D Biol. Crystallogr. 66, 213–221 (2010).

58. Kovach, K. et al. Evolutionary adaptations of biofilms infecting cystic fibrosis lungs promote mechanical toughness by adjusting polysaccharide production. Npj Biofilms Microbiomes 3, 1 (2017).

59. Grant, O. C. et al. Generating 3D Models of Carbohydrates with GLYCAM-Web. bioRxiv 2025.05.08.652828 (2025) doi:10.1101/2025.05.08.652828.

60. Case, D. A. et al. AmberTools. J. Chem. Inf. Model. 63, 6183–6191 (2023).

61. Kirschner, K. N. et al. GLYCAM06: a generalizable biomolecular force field. Carbohydrates. J. Comput. Chem. 29, 622–655 (2008).

62. Tian, C. et al. ff19SB: amino-acid-specific protein backbone parameters trained against quantum mechanics energy surfaces in solution. J. Chem. Theory Comput. 16, 528–552 (2020).

63. Roe, D. R. & Brooks, B. R. A protocol for preparing explicitly solvated systems for stable molecular dynamics simulations. J. Chem. Phys. 153, 054123 (2020).

64. Roe, D. R. & Cheatham, T. E. I. PTRAJ and CPPTRAJ: software for processing and analysis of molecular dynamics trajectory data. J. Chem. Theory Comput. 9, 3084–3095 (2013).

65. Miller, B. R. et al. MMPBSA.py: an efficient program for end-state free energy calculations. J. Chem. Theory Comput. 8, 3314–3321 (2012).

66. Parks, D. H. et al. GTDB: an ongoing census of bacterial and archaeal diversity through a phylogenetically consistent, rank normalized and complete genome-based taxonomy. Nucleic Acids Res. 50, D785–D794 (2022).

67. Seemann, T. Prokka: rapid prokaryotic genome annotation. Bioinforma. Oxf. Engl. 30, 2068–2069 (2014).

68. Parks, D. H., Imelfort, M., Skennerton, C. T., Hugenholtz, P. & Tyson, G. W. CheckM: assessing the quality of microbial genomes recovered from isolates, single cells, and metagenomes. Genome Res. 25, 1043–1055 (2015).

69. Fu, L., Niu, B., Zhu, Z., Wu, S. & Li, W. CD-HIT: accelerated for clustering the next-generation sequencing data. Bioinforma. Oxf. Engl. 28, 3150–3152 (2012).

70. Katoh, K. & Standley, D. M. MAFFT multiple sequence alignment software version 7: improvements in performance and usability. Mol. Biol. Evol. 30, 772–780 (2013).

71. Ashkenazy, H. et al. ConSurf 2016: an improved methodology to estimate and visualize evolutionary conservation in macromolecules. Nucleic Acids Res. 44, W344–350 (2016).

72. Neelamegham, S. et al. Updates to the symbol nomenclature for glycans guidelines. Glycobiology 29, 620–624 (2019).

