## Supplemental Data for "Structure of a key adhesin-exopolysaccharide interaction provides insights into matrix assembly in *Vibrio cholerae* biofilms"

**SUPPLEMENTARY TABLES**

**Table S1. STARANISO scaling information using elliptical truncation of diffraction data.**

Diffraction limits (Å) and corresponding principal axes of the ellipsoid fitted to the diffraction cut-off surface:

| | | | |
|---|---|---|---|
| Diffraction limit #1: | 1.599 | ( 1.0000, 0.0000,0.0000) | ***a**** |
| Diffraction limit #2: | 1.599 | ( 0.0000, 1.0000, 0.0000) | ***b**** |
| Diffraction limit #3: | 1.528 | (0.0000, 0.0000, 1.0000) | ***c**** |

**Criteria used in determination of diffraction limits:**

local($I/\sigma(I)$) >= 1.20

| **Per-reflection cut-off** | **Operational Resolution** |
|---|---|
| $I/\sigma(I)$ >= 2.0 | 1.72 Å for 84,212 reflections |
| $I/\sigma(I)$ >= 1.0 | 1.66 Å for 94,063 reflections |
| $I/\sigma(I)$ >= 0.0 | 1.61 Å for 102,450 reflections |
| All | 1.60 Å for 105,069 reflections |

**Table S2. Data collection and refinement statistics*.**

| | Bap1 + VPS (PDB:10ZX) |
|---|---|
| **DATA COLLECTION** | |
| **Wavelength (Å)** | 0.9793 |
| **Resolution range (Å)** | 76.1 - 1.53 (1.62 - 1.53) |
| **Space group** | $P4_12_12$ |
| **Cell dimensions** $a,b,c,\alpha,\beta,\gamma$ | 71.2 Å, 71.2 Å, 304.5 Å, 90°, 90°, 90° |
| **Total reflections** | 1,977,759 (61,549) |
| **Unique reflections** | 105,069 (5,254) |
| **Multiplicity** | 18.8 (11.7) |
| **Completeness (%)** | 95.0 (51.6) (ellipsoidal) |
| **Mean** $I/\sigma(I)$ | 9.1 (1.6) |
| **Wilson B-factor** | 17.9 |
| $R_{merge}$ | 0.21 (1.25) |
| $R_{pim}$ | 0.05 (0.38) |
| $CC_{1/2}$ | 0.99 (0.66) |
| **REFINEMENT** | |
| **Reflections used in refinement** | 105,064 |
| **Reflections used for** $R_{free}$ | 5,200 |
| $R_{work}$ (%) | 15.4 |
| $R_{free}$ (%) | 17.6 |
| **Number of non-hydrogen atoms** | 5,376 |
| macromolecules | 4,675 |
| ligands | 104 |
| solvent | 597 |
| **Protein residues** | 1,467 |
| **RMSD bond lengths (Å)** | 0.005 |
| **RMSD bond angles (°)** | 0.80 |
| **Ramachandran favored (%)** | 96.5 |
| **Ramachandran allowed (%)** | 3.3 |
| **Ramachandran outliers (%)** | 1.0 |
| **Rotamer outliers (%)** | 0.7 |
| **Molprobity Clashscore** | 0.43 |
| **Average B-factor (Å$^2$)** | 22.5 |
| macromolecules | 20.9 |
| ligands | 28.9 |
| solvent | 33.5 |

*Statistics for the highest-resolution shell are shown in parentheses.

**Table S3. Glycosidic bond dihedral angle comparison.** GW is the GLYCAM-Web model of VPS. RbmB VPS is from PDB:9OZD, chain F. Divergent angles seen in the Bap1 Glc-Glc glycosidic bond are shaded.

| Glycosidic bond dihedral angles ($\Phi$:$O_5$-$C_1$-$O_4$'-$C_4$' $\Psi$: $C_1$-$O_4$'-$C_4$'-$C_3$') | | | | | | | |
|---|---|---|---|---|---|---|---|
| | | | | Δ-Relative to GW | | Δ-Relative to RbmB | |
| **VPS** | **Bond** | $\Phi$ | $\Psi$ | $\lvert\Phi_2-\Phi_1\rvert$ | $\lvert\Psi_2-\Psi_1\rvert$ | $\lvert\Phi_2-\Phi_1\rvert$ | $\lvert\Psi_2-\Psi_1\rvert$ |
| **GW** | L-Gulα(1-4)D-Glcpβ | - 73.2 | 134.2 | 0 | 0 | 4.6 | 8.0 |
| | D-Glcβ(1-4)D-Glcpα | -73.1 | 125.7 | 0 | 0 | 8.7 | 4.5 |
| | D-Glcα(1-4)D-Galpα | 76.0 | -134.5 | 0 | 0 | 1.2 | 5.0 |
| **RbmB** | L-Gulα(1-4)D-Glcpβ | -68.6 | 126.2 | 4.6 | 8.0 | 0 | 0 |
| | D-Glcβ(1-4)D-Glcpα | -81.8 | 130.2 | 8.7 | 4.5 | 0 | 0 |
| | D-Glcα(1-4)D-Galpα | 77.2 | -139.5 | 1.2 | 5.0 | 0 | 0 |
| **Bap1** | *L-Gulα(1-4)D-Glcpβ | -94.3 | 70.7 | 21.1 | 63.5 | 25.7 | 55.5 |
| | D-Glcβ(1-4)D-Glcpα | -103.6 | -64.0 | 30.5 | 170.3 | 21.8 | 165.8 |
| | D-Glcα(1-4)D-Galpα | 83.7 | -110.0 | 7.7 | 24.5 | 6.5 | 29.5 |

* The L-Gulα ring is expected to contain a C4–C5 double bond

**Table S4.** Hydrogen bonding distances observed between Bap1 and VPS during MD simulations of the Bap1-VPS co-complex.

| Ligand (VPS) residue | Protein(Bap1) residue | bond | H-bond distance (Å) in | |
|---|---|---|---|---|
| | | | Crystal structure | MD simulation |
| 1 GulNAcA (α) | 88 Tyr | 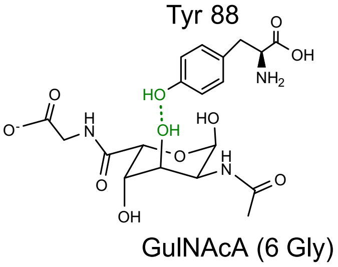<br>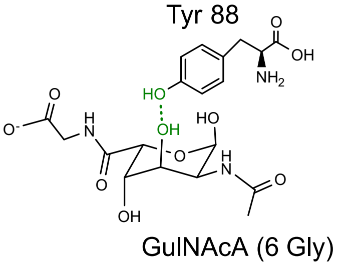 | 3.3 | 4.0 ± 0.7 |
| 1 GulNAcA (α) | 88 Tyr | 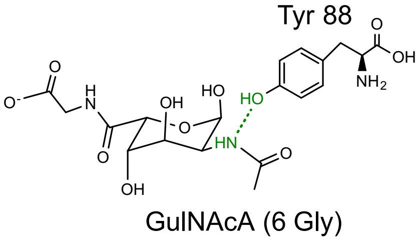<br>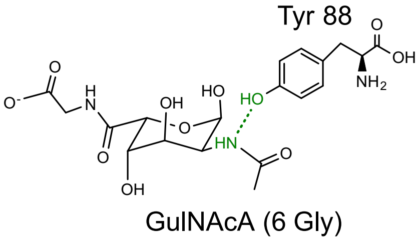 | 2.6 | 4.0 ± 0.5 |
| 1 Gly | 266 Leu | 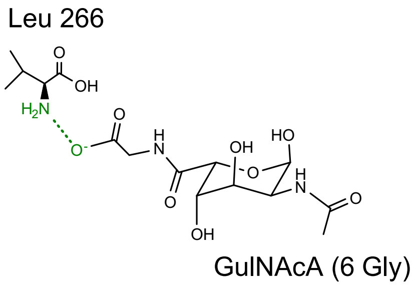<br>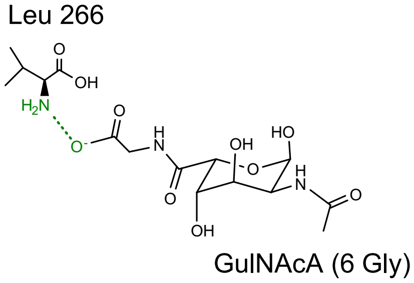 | 2.8 | 5.1 ± 0.5 |

| | | | | |
|---|---|---|---|---|
| 1 Gly | 528 Tyr | 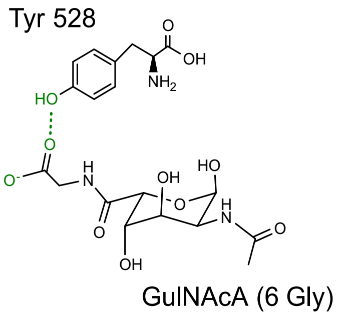  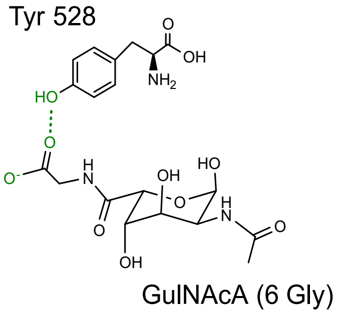  | 2.7 | 4.0 ± 0.6 |
| 2 Glc (β) | 118 Arg | 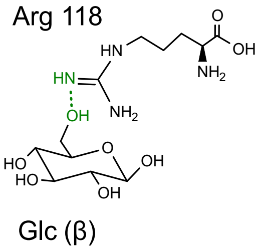  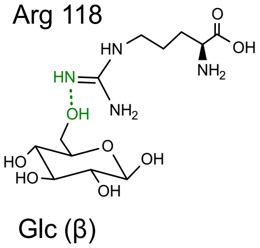  | 2.9 | 2.9 ± 0.0 |
| 2 Glc (β) | 553 Lys | 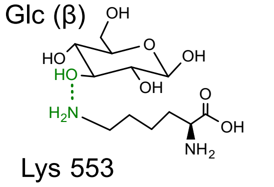  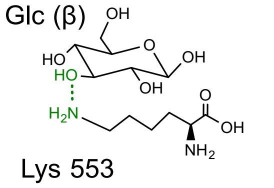  | 2.8 | 4.0 ± 0.3 |
| 3 Glc (α) | 119 Tyr | 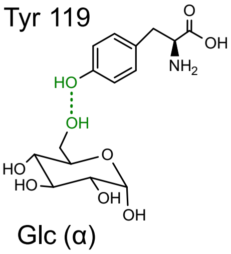  | 2.7 | 3.3 ± 0.1 |

| | | | | |
|---|---|---|---|---|
| | | 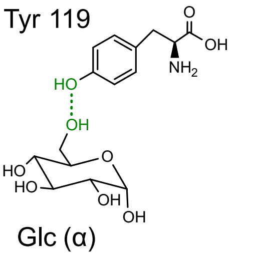 | | |
| 4 Gal (α) | 118 Arg | 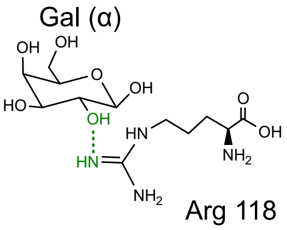 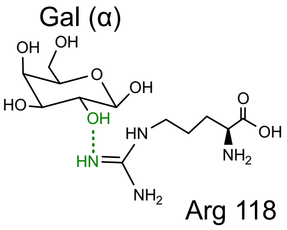 | 3.5 | 4.2 ± 0.2 |
| 4 Gal (α) | 118 Arg | 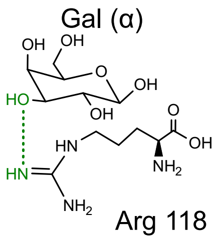 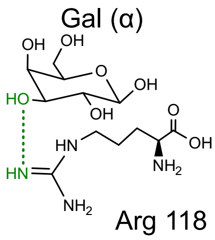 | 2.9 | 3.7 ± 0.1 |

**Table S5.** Ring puckering of the VPS unit observed during MD simulations of the Bap1–VPS co-complex.

a) Bap1–VPS tetrameric fragment co-complex

| Experiment | Ring conformation convergence |
|---|---|
| 1 GulNAcAα | $^{1}C_{4}$ (89.01%) |
| 2 Glcβ | $^{4}C_{1}$ (61.67%) |
| 3 Glcα | $^{4}C_{1}$ (92.23%) |
| 4 Galα | $^{4}C_{1}$ (88.59%) |

b) Bap1–VPS dodecameric fragment co-complex.

| Residue | Ring conformation convergence |
|---|---|
| 1 GulNAcAα | $^{1}C_{4}$ (96.73 %) |
| 2 Glcβ | $^{4}C_{1}$ (69.33%) |
| 3 Glcα | $^{4}C_{1}$ (90.37%) |
| 4 Galα | $^{4}C_{1}$ (95.25%) |
| 5 GulNAcAα | $^{1}C_{4}$ (93.99 %) |
| 6 Glcβ | $^{4}C_{1}$ (67.11%) |
| 7 Glcα | $^{4}C_{1}$ (93.81%) |
| 8 Galα | $^{4}C_{1}$ (94.53%) |
| 9 GulNAcAα | $^{1}C_{4}$ (96.14 %) |
| 10 Glcβ | $^{4}C_{1}$ (78.51%) |
| 11 Glcα | $^{4}C_{1}$ (90.70%) |
| 12 Galα | $^{4}C_{1}$ (94.87%) |

**Table S6.** CHI energy of each glycosidic linkage in the tetrameric fragment of VPS when bound to Bap1 determined from MD simulation.

| | CHI energy of Glycosidic Linkage (kcal/mol) | | |
|---|---|---|---|
| Experiment | GulNAcAα(1→4)Glc | Glcβ(1→4)Glc | Glcα(1→4)Gal |
| run1 | 1.72 | 6.03 | 0.64 |
| run2 | 1.38 | 5.96 | 1.27 |
| run3 | 1.28 | 6.28 | 1.12 |
| Average ± std | 1.46 ± 0.20 | 6.09 ± 0.13 | 1.01 ± 0.27 |

**Table S7.** Contribution of key Bap1 protein residues (contributing >3% of interaction energy) to the formation and stability of Bap1 complexes with tetrameric fragment.

| Wild Type | ΔE Contribution[a] (%) | Point Mutation | ΔE Contribution[a] (%) | p-value[b] |
|---|---|---|---|---|
| W164 | -5.40 ± 0.77 | W164A | -1.39 ± 0.26 | 0.006* |
| R118 | -4.43 ± 0.93 | R118A | -0.20 ± 0.04 | 0.052 |
| D548 | -3.64 ± 0.83 | D548A | -0.10 ± 0.03 | 0.017* |
| Y528 | -3.29 ± 1.40 | Y528A | -0.01 ± 0.00 | 0.056 |
| L598 | -2.79 ± 0.15 | L598A | -0.87 ± 0.06 | 0.001* |
| S265 | -2.58 ± 0.42 | S265A | -1.57 ± 0.41 | 0.038* |
| N356 (Negative Control) | 0.00 ± 0.00 | N356A | 0.00 ± 0.00 | 0.023* |

[a]a negative sign indicates a favorable interaction energy
[b]independent paired t-test
*p-value < 0.05

**Table S8.** Percentage contribution to total interaction energy made by individual ligand residues in the Bap1 complex with a tetrameric VPS fragment.

| Residue | ΔE Contribution[a] (%) |
|---|---|
| 1 GulNAcAα | -14.7 ± 2.0 |
| 1 Gly | 3.6 ± 0.9 |
| 2 Glcβ | -4.5 ± 0.9 |
| 3 Glcα | -3.8 ± 0.4 |
| 4 Galα | -1.0 ± 0.2 |

[a]a negative sign indicates a favorable interaction energy

**Table S9.** Primers used in this study.

| Primer Name | Primer Sequence (5' to 3')* | Description |
|---|---|---|
| **Mutant constructs** | | |
| ZJ-P-019 | CTTGTCATCGTCATCCTTGTAATCGATA | Universal sequencer for 3xFLAG |
| ZJ-P-020 | TGCAATTGGTGCTAATAGCCAAAAC | *bap1* sequencing Forward 2 |
| ZJ-P-026 | AGCAGCATTTTGAAAACTTCCGC | 2.7kb upstream of *bap1* |
| ZJ-P-027 | ATGAAATTCACGATAACCAGAAAACCG | 2.7kb downstream of *bap1* |
| XH-P-28 | GTGGTGAAACCCACTATCTGGATC | 0.5kb upstream of *bap1* |
| XH-P-29 | TCTTGCTGGCTAAACAACGCAT | 0.5kb downstream of *bap1* |
| JY-P-140 | AATCAAACCGGGCTTTAAATTTCATCTCGAC | 3kb upstream of *bap1* |
| JY-P-141 | CATGATATGCAACATCTACTGAAAGAGGTGCA | 3kb downstream of *bap1* |
| BN-P-01 | CACCTTTGAAGGTAATAAGTTCGCGAATGGTGGTTATATCC | *bap1* Y88F Forward |
| BN-P-02 | GGATATAACCACCATTCGCGAACTTATTACCTTCAAAGGTG | *bap1* Y88F Reverse |
| BN-P-03 | GAGGCGTGATTGCCGATGCTATGTATGCACCAGCAGCCGCG G | *bap1* R118M Forward |
| BN-P-04 | CCGCGGCTGCTGGTGCATACATAGCATCGGCAATCACGCCT C | *bap1* R118M Reverse |
| BN-P-05 | GCTGAAATCTGCTTCAGGCGCGCGTTCTGTAGGTGATATTG | *bap1* W164A Forward |
| BN-P-06 | CAATATCACCTACAGAACGCGCGCCTGAAGCAGATTTCAGC | *bap1* W164A Reverse |
| BN-P-07 | GGCTAACGATGACCTGAACAAAGGCAAAATTGGTGTCTCCG C | *bap1* S551K Forward |
| BN-P-08 | GCGGAGACACCAATTTTGCCTTTGTTCAGGTCATCGTTAGCC | *bap1* S551K Reverse |
| BN-P-09 | GTGGCTAACGATGACCTGAACGCCGGCAAAATTGGTGTCTC CG | *bap1* S551A Forward |
| BN-P-10 | CGGAGACACCAATTTTGCCGGCGTTCAGGTCATCGTTAGCC | *bap1* S551A |

| | | |
|---|---|---|
| | AC | Reverse |
| BN-P-11 | CGATGACCTGAACAGCGGCGCAATTGGTGTCTCCGCTTACG | *bap1* K553A Forward |
| BN-P-12 | CGTAAGCGGAGACACCAATTGCGCCGCTGTTCAGGTCATCG | *bap1* K553A Reverse |
| BN-P-13 | GCGAACTCTTCTGGCACCAAGTGGGAATACCCAGTCGTGG | *bap1* L598K Forward |
| BN-P-14 | CCACGACTGGGTATTCCCACTTGGTGCCAGAAGAGTTCGC | *bap1* L598K Reverse |
| BN-P-19 | CACCTTTGAAGGTAATAAGTGGGCGAATGGTGGTTATATC | *bap1* Y88W Forward |
| BN-P-20 | GATATAACCACCATTCGCCCACTTATTACCTTCAAAGGTG | *bap1* Y88W Reverse |
| BN-P-21 | GTGATTGCCGATGCTGCCTATGCACCAGC | *bap1* R118A Forward |
| BN-P-22 | GCTGGTGCATAGGCAGCATCGGCAATCAC | *bap1* R118A Reverse |
| BN-P-29 | CTAACGATGACCTGGCCAGCGGCAAAATTG | *bap1* N550A Forward |
| BN-P-30 | CAATTTTGCCGCTGGCCAGGTCATCGTTAG | *bap1* N550A Reverse |
| Y88A-F | GTCACCTTTGAAGGTAATAAGGCCGCGAATGGTGGTTATATC CGC | *bap1* Y88A Forward |
| Y88A-R | GCGGATATAACCACCATTCGCGGCCTTATTACCTTCAAAGGT GAC | *bap1* Y88A Reverse |
| Y88F-F | GTCACCTTTGAAGGTAATAAGTTCGCGAATGGTGGTTATATC CGC | *bap1* Y88F Forward |
| Y88F-R | GCGGATATAACCACCATTCGCGAACTTATTACCTTCAAAGGT GAC | *bap1* Y88F Reverse |
| Y88W-F | GTCACCTTTGAAGGTAATAAGTGGGCGAATGGTGGTTATATC GCGCCC | *bap1* Y88W Forward |
| Y88W-R | GGGCGCGGATATAACCACCATTCGCCCACTTATTACCTTCAA AGGTGAC | *bap1* Y88W Reverse |
| R118A-F | GGAGGCGTGATTGCCGATGCTGCCTATGCACCAGCAGCCGC GGATC | *bap1* R118A Forward |
| R118A-R | GATCCGCGGCTGCTGGTGCATAGGCAGCATCGGCAATCACG CCTCC | *bap1* R118A Reverse |
| R118M-F | GGAGGCGTGATTGCCGATGCTATGTATGCACCAGCAGCCGC GGAT | *bap1* R118M Forward |
| R118M-R | GATCCGCGGCTGCTGGTGCATACATAGCATCGGCAATCACG CCTCC | *bap1* R118M Reverse |
| Y119F-F | GGCGTGATTGCCGATGCTCGCTTCGCACCAGCAGCCGCGGA TCTC | *bap1* Y119F Forward |
| Y119F-R | GAGATCCGCGGCTGCTGGTGCGAAGCGAGCATCGGCAATCA CGCC | *bap1* Y119F Reverse |

| | | |
|---|---|---|
| R165A-F | CTGAAATCTGCTTCAGGCTGGGCCTCTGTAGGTGATATTGCACTGG | *bap1* R165A Forward |
| R165A-R | CCAGTGCAATATCACCTACAGAGGCCCAGCCTGAAGCAGATTTCAG | *bap1* R165A Reverse |
| Y528F-F | CATGGGCATTGTGTACGCGGGTTTTTACGCGGTTGATATGTACGATGCGC | *bap1* Y528F Forward |
| Y528F-R | GCGCATCGTACATATCAACCGCGTAAAAACCCGCGTACACAATGCCCATG | *bap1* Y528F Reverse |
| N550A-F | GTCTGTGGCTAACGATGACCTGGCCAGCGGCAAAATTGGTGTCTCC | *bap1* N550A Forward |
| N550A-R | GGAGACACCAATTTTGCCGCTGGCCAGGTCATCGTTAGCCACAGAC | *bap1* N550A Reverse |
| S551A-F | GTGGCTAACGATGACCTGAACGCGGGCAAAATTGGTGTCTCCGCTTAC | *bap1* S551A Forward |
| S551A-R | GTAAGCGGAGACACCAATTTTGCCCGCGTTCAGGTCATCGTTAGCCAC | *bap1* S551A Reverse |
| S551K-F | CTGTGGCTAACGATGACCTGAACAAAGGCAAAATTGGTGTCTCCGCTTAC | *bap1* S551K Forward |
| S551K-R | GTAAGCGGAGACACCAATTTTGCCTTTGTTCAGGTCATCGTTAGCCACAG | *bap1* S551K Reverse |
| K553A-F | CTAACGATGACCTGAACAGCGGCGCAATTGGTGTCTCCGCTTACGAC | *bap1* K553A Forward |
| K553A-R | GTCGTAAGCGGAGACACCAATTGCGCCGCTGTTCAGGTCATCGTTAG | *bap1* K553A Reverse |
| T597A-F | CATTATTGCGAACTCTTCTGGCGCCTTGTGGGAATACCCAGTCGTG | *bap1* T597A Forward |
| T597A-R | CACGACTGGGTATTCCCACAAGGCGCCAGAAGAGTTCGCAATAATG | *bap1* T597A Reverse |
| L598K-F | CATTATTGCGAACTCTTCTGGCACCAAATGGGAATACCCAGTCGTGGCAG | *bap1* L598K Forward |
| L598K-R | CTGCCACGACTGGGTATTCCCATTTGGTGCCAGAAGAGTTCGCAATAATG | *bap1* L598K Reverse |

*lowercase nucleotides indicate overlapping sequence for SOE or nucleotide change for aa mutations

**Table S10.** Strains used in this study.

| Strain Name in Manuscript | Genotype and Antibiotic Resistance | Description | Strain# & Reference |
|---|---|---|---|
| WT background | C6706, $Sm^R$ | A streptomycin-resistant variant of the WT O1 El Tor biotype C6706str2. Serves as the background for all strains reported in this manuscript | JY016[1] |
| Rg background | *vpvC*$^{W240R}$, $Sm^R$ | Missense mutation in the *Vibrio cholerae* O1 El tor strain that elevates the level of cyclic-di-GMP. Rugose phenotype. Serves as the parental strain for all mutants in this manuscript | JY028[2] |
| **Strains Used for Biofilm Morphology and Adhesion Assays** | | | |
| Rg | *vpvC*$^{W240R}$, Δ*VC1807*::$P_{tac}$-*mNeonGreen*, $Spec^R$ | Rugose strain with mNeonGreen fluorescent protein inserted at a non-essential site in the genome (VC1807) | JY451[3] |
| Rg Δ*bap1* Δ*rbmC* | *vpvC*$^{W240R}$, Δ*bap1*, Δ*rbmC* Δ*VC1807*::$P_{tac}$-*mNeonGreen*, $Spec^R$ | Rugose strain with mNeonGreen fluorescent protein inserted at a non-essential site in the genome (VC1807) and *rbmC* and *bap1* deletion | JY458, This Study |
| Δ*bap1* | *vpvC*$^{W240R}$, Δ*bap1*, Δ*VC1807*::$P_{tac}$-*mNeonGreen*, $Spec^R$ | Clean deletion of *bap1* by cotransformation with Δ*VC1807*::$P_{tac}$-*mNeonGreen*, $Spec^R$ | ZJ053, This Study |
| Δ*rbmC* | *vpvC*$^{W240R}$, Δ*rbmC*, Δ*VC1807*::$P_{tac}$-*mNeonGreen*, $Spec^R$ | Clean deletion of *rbmC* by cotransformation with Δ*VC1807*::$P_{tac}$-*mNeonGreen*, $Spec^R$ | ZJ033, This Study |
| Δ*bap1* Δ*rbmC* | *vpvC*$^{W240R}$, Δ*bap1*, Δ*rbmC* | Rugose strain with *rbmC* and *bap1* deletion | JY301, This Study |
| *bap1*$^{Y88F}$ | *vpvC*$^{W240R}$, Δ*rbmC*, *bap1*$^{Y88F}$, Δ*VC1807*::$P_{tac}$-*mNeonGreen*, $Spec^R$ | Introduction of mutated *bap1* to Δ*bap1* Δ*rbmC* by cotransformation with Δ*VC1807*::$P_{tac}$-*mNeonGreen*, $Spec^R$ | BN11, This Study |
| *bap1*$^{W164A}$ | *vpvC*$^{W240R}$, Δ*rbmC*, *bap1*$^{W164A}$, Δ*VC1807*::$P_{tac}$-*mNeonGreen*, $Spec^R$ | Introduction of mutated *bap1* to Δ*bap1* Δ*rbmC* by cotransformation with Δ*VC1807*::$P_{tac}$-*mNeonGreen*, $Spec^R$ | BN12, This Study |

| | | | |
|---|---|---|---|
| *bap1*$^{Y88W}$ | *vpvC*$^{W240R}$, Δ*rbmC*, *bap1*$^{Y88W}$, Δ*VC1807*::*P*$_{tac}$-*mNeonGreen*, Spec$^{R}$ | Introduction of mutated *bap1* to Δ*bap1* Δ*rbmC* by cotransformation with Δ*VC1807*::*P*$_{tac}$-*mNeonGreen*, Spec$^{R}$ | BN14, This Study |
| *bap1*$^{R118A, L598K}$ | *vpvC*$^{W240R}$, Δ*rbmC*, *bap1*$^{R118A, L598K}$, Δ*VC1807*::*P*$_{tac}$-*mNeonGreen*, Spec$^{R}$ | Introduction of mutated *bap1* to Δ*bap1* Δ*rbmC* by cotransformation with Δ*VC1807*::*P*$_{tac}$-*mNeonGreen*, Spec$^{R}$ | BN15, This Study |
| *bap1*$^{K553A}$ | *vpvC*$^{W240R}$, Δ*rbmC*, *bap1*$^{K553A}$, Δ*VC1807*::*P*$_{tac}$-*mNeonGreen*, Spec$^{R}$ | Introduction of mutated *bap1* to Δ*bap1* Δ*rbmC* by cotransformation with Δ*VC1807*::*P*$_{tac}$-*mNeonGreen*, Spec$^{R}$ | BN16, This Study |
| *bap1*$^{S551K}$ | *vpvC*$^{W240R}$, Δ*rbmC*, *bap1*$^{S551K}$, Δ*VC1807*::*P*$_{tac}$-*mNeonGreen*, Spec$^{R}$ | Introduction of mutated *bap1* to Δ*bap1* Δ*rbmC* by cotransformation with Δ*VC1807*::*P*$_{tac}$-*mNeonGreen*, Spec$^{R}$ | BN17, This Study |
| *bap1*$^{L598K}$ | *vpvC*$^{W240R}$, Δ*rbmC*, *bap1*$^{L598K}$, Δ*VC1807*::*P*$_{tac}$-*mNeonGreen*, Spec$^{R}$ | Introduction of mutated *bap1* to Δ*bap1* Δ*rbmC* by cotransformation with Δ*VC1807*::*P*$_{tac}$-*mNeonGreen*, Spec$^{R}$ | BN18, This Study |
| *bap1*$^{S551A}$ | *vpvC*$^{W240R}$, Δ*rbmC*, *bap1*$^{S551A}$, Δ*VC1807*::*P*$_{tac}$-*mNeonGreen*, Spec$^{R}$ | Introduction of mutated *bap1* to Δ*bap1* Δ*rbmC* by cotransformation with Δ*VC1807*::*P*$_{tac}$-*mNeonGreen*, Spec$^{R}$ | BN19, This Study |
| *bap1*$^{R118A}$ | *vpvC*$^{W240R}$, Δ*rbmC*, *bap1*$^{R118A}$, Δ*VC1807*::*P*$_{tac}$-*mNeonGreen*, Spec$^{R}$ | Introduction of mutated *bap1* to Δ*bap1* Δ*rbmC* by cotransformation with Δ*VC1807*::*P*$_{tac}$-*mNeonGreen*, Spec$^{R}$ | BN20, This Study |
| *bap1*$^{N550A}$ | *vpvC*$^{W240R}$, Δ*rbmC*, *bap1*$^{N550A}$, Δ*VC1807*::*P*$_{tac}$-*mNeonGreen*, Spec$^{R}$ | Introduction of mutated *bap1* to Δ*bap1* Δ*rbmC* by cotransformation with Δ*VC1807*::*P*$_{tac}$-*mNeonGreen*, Spec$^{R}$ | BN21, This Study |

| | | | |
|---|---|---|---|
| *bap1*$^{R118M}$ | *vpvC*$^{W240R}$, Δ*rbmC*, *bap1*$^{R118M}$, Δ*VC1807*::*P*$_{tac}$-*mNeonGreen*, Spec$^{R}$ | Introduction of mutated *bap1* to Δ*bap1* Δ*rbmC* by cotransformation with Δ*VC1807*::*P*$_{tac}$-*mNeonGreen*, Spec$^{R}$ | BN27, This Study |
| *bap1*$^{T597A}$ | *vpvC*$^{W240R}$, Δ*rbmC*, *bap1*$^{T597A}$, Δ*VC1807*::*P*$_{tac}$-*mNeonGreen*, Spec$^{R}$ | Introduction of mutated *bap1* to Δ*bap1* Δ*rbmC* by cotransformation with Δ*VC1807*::*P*$_{tac}$-*mNeonGreen*, Spec$^{R}$ | JC01, This Study |
| *bap1*$^{Y88A}$ | *vpvC*$^{W240R}$, Δ*rbmC*, *bap1*$^{Y88A}$, Δ*VC1807*::*P*$_{tac}$-*mNeonGreen*, Spec$^{R}$ | Introduction of mutated *bap1* to Δ*bap1* Δ*rbmC* by cotransformation with Δ*VC1807*::*P*$_{tac}$-*mNeonGreen*, Spec$^{R}$ | JC02, This Study |
| **Strains Used for *in situ* Staining** | | | |
| *bap1-3XFLAG* | *vpvC*$^{W240R}$, *bap1-3XFLAG*, Δ*VC1807*::*P*$_{tac}$-*mNeonGreen*, Spec$^{R}$ | 3XFLAG tagged Bap1 | JY488[3] |
| Δ*rbmC* | *vpvC*$^{W240R}$, Δ*rbmC*, Δ*VC1807*::*P*$_{tac}$-*mNeonGreen*, Spec$^{R}$ | Non-tagged Bap1 with *rbmC* deleted | ZJ033, This Study |
| Δ*rbmC bap1-3XFLAG* | *vpvC*$^{W240R}$, Δ*rbmC*, *bap1-3XFLAG*, Δ*VC1807*::*P*$_{tac}$-*mNeonGreen*, Spec$^{R}$ | 3XFLAG tagged Bap1 with *rbmC* deleted | ZJ065, This Study |
| *bap1*$^{Y88F}$-*3XFLAG* | *vpvC*$^{W240R}$, Δ*rbmC*, *bap1*$^{Y88F}$-*3XFLAG*, Δ*VC1807*::*P*$_{tac}$-*mNeonGreen*, Kan$^{R}$ | 3XFLAG tagged Bap1$^{Y88F}$ with *rbmC* deleted | BN22, This Study |
| *bap1*$^{W164A}$-*3XFLAG* | *vpvC*$^{W240R}$, Δ*rbmC*, *bap1*$^{W164A}$-*3XFLAG*, Δ*VC1807*::*P*$_{tac}$-*mNeonGreen*, Kan$^{R}$ | 3XFLAG tagged Bap1$^{W164A}$ with *rbmC* deleted | BN23, This Study |
| *bap1*$^{Y88W}$-*3XFLAG* | *vpvC*$^{W240R}$, Δ*rbmC*, *bap1*$^{Y88W}$-*3XFLAG*, Δ*VC1807*::*P*$_{tac}$-*mNeonGreen*, Kan$^{R}$ | 3XFLAG tagged Bap1$^{Y88W}$ with *rbmC* deleted | BN24, This Study |
| *bap1*$^{K553A}$-*3XFLAG* | *vpvC*$^{W240R}$, Δ*rbmC*, *bap1*$^{K553A}$-*3XFLAG*, Δ*VC1807*::*P*$_{tac}$-*mNeonGreen*, Kan$^{R}$ | 3XFLAG tagged Bap1$^{K553A}$ with *rbmC* deleted | BN25, This Study |

| | | | |
|---|---|---|---|
| *bap1*$^{S551K}$-*3XFLAG* | *vpvC*$^{W240R}$, Δ*rbmC*, *bap1*$^{S551K}$-*3XFLAG*, Δ*VC1807*::*P*$_{tac}$-*mNeonGreen*, Kan$^R$ | 3XFLAG tagged Bap1$^{S551K}$ with *rbmC* deleted | BN26, This Study |
| *bap1*$^{L598K}$-*3XFLAG* | *vpvC*$^{W240R}$, Δ*rbmC*, *bap1*$^{L598K}$-*3XFLAG*, Δ*VC1807*::*P*$_{tac}$-*mNeonGreen*, Kan$^R$ | 3XFLAG tagged Bap1$^{L598K}$ with *rbmC* deleted | BN28, This Study |
| *bap1*$^{S551A}$-*3XFLAG* | *vpvC*$^{W240R}$, Δ*rbmC*, *bap1*$^{S551A}$-*3XFLAG*, Δ*VC1807*::*P*$_{tac}$-*mNeonGreen*, Kan$^R$ | 3XFLAG tagged Bap1$^{S551A}$ with *rbmC* deleted | BN29, This Study |
| *bap1*$^{R118A}$-*3XFLAG* | *vpvC*$^{W240R}$, Δ*rbmC*, *bap1*$^{R118A}$-*3XFLAG*, Δ*VC1807*::*P*$_{tac}$-*mNeonGreen*, Kan$^R$ | 3XFLAG tagged Bap1$^{R118A}$ with *rbmC* deleted | BN30, This Study |
| *bap1*$^{N550A}$-*3XFLAG* | *vpvC*$^{W240R}$, Δ*rbmC*, *bap1*$^{N550A}$-*3XFLAG*, Δ*VC1807*::*P*$_{tac}$-*mNeonGreen*, Kan$^R$ | 3XFLAG tagged Bap1$^{N550A}$ with *rbmC* deleted | BN31, This Study |
| *bap1*$^{R118A, L598K}$-*3XFLAG* | *vpvC*$^{W240R}$, Δ*rbmC*, *bap1*$^{R118A, L598K}$-*3XFLAG*, Δ*VC1807*::*P*$_{tac}$-*mNeonGreen*, Kan$^R$ | 3XFLAG tagged Bap1$^{R118A, L598K}$ with *rbmC* deleted | BN32, This Study |
| *bap1*$^{R118M}$-*3XFLAG* | *vpvC*$^{W240R}$, Δ*rbmC*, *bap1*$^{R118M}$-*3XFLAG*, Δ*VC1807*::*P*$_{tac}$-*mNeonGreen*, Kan$^R$ | 3XFLAG tagged Bap1$^{R118M}$ with *rbmC* deleted | BN33, This Study |
| *bap1*$^{T597A}$-*3XFLAG* | *vpvC*$^{W240R}$, Δ*rbmC*, *bap1*$^{T597A}$-*3XFLAG*, Δ*VC1807*::*P*$_{tac}$-*mNeonGreen*, Kan$^R$ | 3XFLAG tagged Bap1$^{T597A}$ with *rbmC* deleted | JC05, This Study |
| *bap1*$^{Y88A}$-*3XFLAG* | *vpvC*$^{W240R}$, Δ*rbmC*, *bap1*$^{Y88A}$-*3XFLAG*, Δ*VC1807*::*P*$_{tac}$-*mNeonGreen*, Kan$^R$ | 3XFLAG tagged Bap1$^{Y88A}$ with *rbmC* deleted | JC06, This Study |
| **Strains used for Complementation Assays** | | | |
| *bap1*$^{S551K}$ +*P*$_{BAD}$-*bap1* | *pEVS* (*P*$_{BAD}$_*bap1*, Kan$^R$), *vpvC*$^{W240R}$, Δ*rbmC*, *bap1*$^{S551K}$, Δ*VC1807*::*P*$_{tac}$-*mNeonGreen*, Spec$^R$ | *pEVS* containing *bap1* with an arabinose inducible promoter was mated into the indicated strain for complementation | BN34, This Study |
| *bap1*$^{L598K}$ +*P*$_{BAD}$-*bap1* | *pEVS* (*P*$_{BAD}$_*bap1*, Kan$^R$), *vpvC*$^{W240R}$, Δ*rbmC*, *bap1*$^{L598K}$, Δ*VC1807*::*P*$_{tac}$-*mNeonGreen*, Spec$^R$ | *pEVS* containing *bap1* with an arabinose inducible promoter was mated into the indicated strain for complementation | BN35, This Study |

| | | | |
|---|---|---|---|
| *bap1*$^{S551A}$ +$P_{BAD}$-*bap1* | *pEVS* ($P_{BAD}$_*bap1*, Kan$^{R}$), *vpvC*$^{W240R}$, Δ*rbmC, bap1*$^{S551A}$, Δ*VC1807*::$P_{tac}$-*mNeonGreen*, Spec$^{R}$ | *pEVS* containing *bap1* with an arabinose inducible promoter was mated into the indicated strain for complementation | BN36, This Study |
| *bap1*$^{N550A}$ +$P_{BAD}$-*bap1* | *pEVS* ($P_{BAD}$_*bap1*, Kan$^{R}$), *vpvC*$^{W240R}$, Δ*rbmC, bap1*$^{N550A}$, Δ*VC1807*::$P_{tac}$-*mNeonGreen*, Spec$^{R}$ | *pEVS* containing *bap1* with an arabinose inducible promoter was mated into the indicated strain for complementation | BN37, This Study |
| *bap1*$^{R118, L598K}$ +$P_{BAD}$-*bap1* | *pEVS* ($P_{BAD}$_*bap1*, Kan$^{R}$), *vpvC*$^{W240R}$, Δ*rbmC*, *bap1*$^{R118A, L598K}$, Δ*VC1807*::$P_{tac}$-*mNeonGreen*, Spec$^{R}$ | *pEVS* containing *bap1* with an arabinose inducible promoter was mated into the indicated strain for complementation | BN38, This Study |
| **Strains Used for Rheological Measurement** | | | |
| Δ*pomA* | *vpvC*$^{W240R}$, Δ*rbmC*, Δ*pomA*, Δ*VC1807*::$P_{tac}$-*mNeonGreen*, Kan$^{R}$ | Deletion of *pomA* by cotransformation ZJ033 with Δ*VC1807*::$P_{tac}$-*mNeonGreen*, Kan$^{R}$ | JC15, This Study |
| Δ*pomA*, *bap1*$^{L598K}$ | *vpvC*$^{W240R}$, Δ*rbmC*, Δ*pomA*, *bap1*$^{L598K}$, Δ*VC1807*::$P_{tac}$-*mNeonGreen*, Kan$^{R}$ | Deletion of *pomA* by cotransformation BN18 with Δ*VC1807*::$P_{tac}$-*mNeonGreen*, Kan$^{R}$ | JC07, This Study |
| Δ*pomA*, *bap1*$^{S551K}$ | *vpvC*$^{W240R}$, Δ*rbmC*, Δ*pomA*, *bap1*$^{S551K}$, Δ*VC1807*::$P_{tac}$-*mNeonGreen*, Kan$^{R}$ | Deletion of *pomA* by cotransformation BN17 with Δ*VC1807*::$P_{tac}$-*mNeonGreen*, Kan$^{R}$ | JC08, This Study |
| Δ*pomA*, *bap1*$^{S551A}$ | *vpvC*$^{W240R}$, Δ*rbmC*, Δ*pomA*, *bap1*$^{S551A}$, Δ*VC1807*::$P_{tac}$-*mNeonGreen*, Kan$^{R}$ | Deletion of *pomA* by cotransformation BN19 with Δ*VC1807*::$P_{tac}$-*mNeonGreen*, Kan$^{R}$ | JC09, This Study |
| Δ*pomA*, *bap1*$^{R118A, L598K}$ | *vpvC*$^{W240R}$, Δ*rbmC*, Δ*pomA*, *bap1*$^{R118A, L598K}$, Δ*VC1807*::$P_{tac}$-*mNeonGreen*, Kan$^{R}$ | Deletion of *pomA* by cotransformation BN15 with Δ*VC1807*::$P_{tac}$-*mNeonGreen*, Kan$^{R}$ | JC10, This Study |
| **Strains Used for Western Blots** | | | |
| Δ*vpsL bap1-3XFLAG* | *vpvC*$^{W240R}$, Δ*vpsL*, *bap1-3XFLAG*, Δ*VC1807*::$P_{tac}$- | *vpsL* was deleted in the indicated strain for Western Blots | ZJ063[4] |

| | | | |
|---|---|---|---|
| | *mNeonGreen,* Spec$^{R}$ | | |
| Δ*vpsL* Δ*rbmC bap1*$^{L598K}$*-3XFLAG* | *vpvC*$^{W240R}$*,* Δ*vpsL,* Δ*rbmC, bap1*$^{L598K}$*-3XFLAG,* Δ*VC1807::P*$_{tac}$*-mNeonGreen,* Spec$^{R}$ | *vpsL* was deleted in the indicated strain for Western Blots | JC11 |
| Δ*vpsL* Δ*rbmC bap1*$^{S551K}$*-3XFLAG* | *vpvC*$^{W240R}$*,* Δ*vpsL,* Δ*rbmC, bap1*$^{S551K}$*-3XFLAG,* Δ*VC1807::P*$_{tac}$*-mNeonGreen,* Spec$^{R}$ | *vpsL* was deleted in the indicated strain for Western Blots | JC12 |
| Δ*vpsL* Δ*rbmC bap1*$^{S551A}$*-3XFLAG* | *vpvC*$^{W240R}$*,* Δ*vpsL,* Δ*rbmC, bap1*$^{S551A}$*-3XFLAG,* Δ*VC1807::P*$_{tac}$*-mNeonGreen,* Spec$^{R}$ | *vpsL* was deleted in the indicated strain for Western Blots | JC13 |
| Δ*vpsL* Δ*rbmC bap1*$^{L598K, R118A}$*-3XFLAG* | *vpvC*$^{W240R}$*,* Δ*vpsL,* Δ*rbmC, bap1*$^{L598K, R118A}$*-3XFLAG,* Δ*VC1807::P*$_{tac}$*-mNeonGreen,* Spec$^{R}$ | *vpsL* was deleted in the indicated strain for Western Blots | JC14 |
| Δ*vpsL* Δ*rbmC bap1*$_{\Delta\beta\text{-propeller}}$ | *vpvC*$^{W240R}$*,* Δ*vpsL,* Δ*rbmC, bap1*$_{\Delta\beta\text{-propeller}}$*,* Δ*VC1807::P*$_{tac}$*-mNeonGreen,* Kan$^{R}$ | Negative control for Western Blots | XH006[4] |

| **Strain Name in Manuscript** | **Genotype and Antibiotic Resistance** | **Description** | **Strain# & Reference** |
|---|---|---|---|
| WT background | C6706, Sm$^{R}$ | A streptomycin-resistant variant of the WT O1 El Tor biotype C6706str2. Serves as the background for all strains reported in this manuscript | JY016[1] |
| Rg background | *vpvC*$^{W240R}$, Sm$^{R}$ | Missense mutation in the *Vibrio cholerae* O1 El tor strain that elevates the level of cyclic-di-GMP. Rugose phenotype. Serves as the parental strain for most of the mutants in this manuscript | JY028[2] |
| **Strains Used for Biofilm Morphology and Adhesion Assays** | | | |

| | | | |
|---|---|---|---|
| Rg | *vpvC*$^{W240R}$, Δ*VC1807*::*P*$_{tac}$-*mNeonGreen*, Spec$^R$ | Rugose strain with mNeonGreen fluorescent protein inserted at a non-essential site in the genome (VC1807) | JY451[3] |
| Rg Δ*bap1* Δ*rbmC* | *vpvC*$^{W240R}$, Δ*bap1*, Δ*rbmC* Δ*VC1807*::*P*$_{tac}$-*mNeonGreen*, Spec$^R$ | Rugose strain with mNeonGreen fluorescent protein inserted at a non-essential site in the genome (VC1807) and *rbmC* and *bap1* deletion | JY458, This Study |
| Δ*bap1* | *vpvC*$^{W240R}$, Δ*bap1*, Δ*VC1807*::*P*$_{tac}$-*mNeonGreen*, Spec$^R$ | Clean deletion of *bap1* by cotransformation with Δ*VC1807*::*P*$_{tac}$-*mNeonGreen*, Spec$^R$ | ZJ053, This Study |
| Δ*rbmC* | *vpvC*$^{W240R}$, Δ*rbmC*, Δ*VC1807*::*P*$_{tac}$-*mNeonGreen*, Spec$^R$ | Clean deletion of *rbmC* by cotransformation with Δ*VC1807*::*P*$_{tac}$-*mNeonGreen*, Spec$^R$ | ZJ033, This Study |
| Δ*bap1* Δ*rbmC* | *vpvC*$^{W240R}$, Δ*bap1*, Δ*rbmC* | Rugose strain with *rbmC* and *bap1* deletion | JY301, This Study |
| *bap1*$^{Y88F}$ | *vpvC*$^{W240R}$, Δ*rbmC*, *bap1*$^{Y88F}$, Δ*VC1807*::*P*$_{tac}$-*mNeonGreen*, *Spec*$^R$ | Introduction of mutated *bap1* to Δ*bap1* Δ*rbmC* by cotransformation with Δ*VC1807*::*P*$_{tac}$-*mNeonGreen*, Spec$^R$ | BN11, This Study |
| *bap1*$^{W164A}$ | *vpvC*$^{W240R}$, Δ*rbmC*, *bap1*$^{W164A}$, Δ*VC1807*::*P*$_{tac}$-*mNeonGreen*, *Spec*$^R$ | Introduction of mutated *bap1* to Δ*bap1* Δ*rbmC* by cotransformation with Δ*VC1807*::*P*$_{tac}$-*mNeonGreen*, Spec$^R$ | BN12, This Study |
| *bap1*$^{Y88W}$ | *vpvC*$^{W240R}$, Δ*rbmC*, *bap1*$^{Y88W}$, Δ*VC1807*::*P*$_{tac}$-*mNeonGreen*, *Spec*$^R$ | Introduction of mutated *bap1* to Δ*bap1* Δ*rbmC* by cotransformation with Δ*VC1807*::*P*$_{tac}$-*mNeonGreen*, Spec$^R$ | BN14, This Study |
| *bap1*$^{R118A, L598K}$ | *vpvC*$^{W240R}$, Δ*rbmC*, *bap1*$^{R118A, L598K}$, Δ*VC1807*::*P*$_{tac}$-*mNeonGreen*, *Spec*$^R$ | Introduction of mutated *bap1* to Δ*bap1* Δ*rbmC* by cotransformation with Δ*VC1807*::*P*$_{tac}$-*mNeonGreen*, Spec$^R$ | BN15, This Study |

| | | | |
|---|---|---|---|
| *$bap1^{K553A}$* | *$vpvC^{\mathrm{W240R}}$, ΔrbmC, $bap1^{K553A}$, ΔVC1807::$P_{tac}$-mNeonGreen, $Spec^{\mathrm{R}}$* | Introduction of mutated *bap1* to Δ*bap1* Δ*rbmC* by cotransformation with Δ*VC1807*::*$P_{tac}$-mNeonGreen,* $\mathrm{Spec}^{\mathrm{R}}$ | BN16, This Study |
| *$bap1^{S551K}$* | *$vpvC^{\mathrm{W240R}}$, ΔrbmC, $bap1^{S551K}$, ΔVC1807::$P_{tac}$-mNeonGreen, $Spec^{\mathrm{R}}$* | Introduction of mutated *bap1* to Δ*bap1* Δ*rbmC* by cotransformation with Δ*VC1807*::*$P_{tac}$-mNeonGreen,* $\mathrm{Spec}^{\mathrm{R}}$ | BN17, This Study |
| *$bap1^{L598K}$* | *$vpvC^{\mathrm{W240R}}$, ΔrbmC, $bap1^{L598K}$, ΔVC1807::$P_{tac}$-mNeonGreen, $Spec^{\mathrm{R}}$* | Introduction of mutated *bap1* to Δ*bap1* Δ*rbmC* by cotransformation with Δ*VC1807*::*$P_{tac}$-mNeonGreen,* $\mathrm{Spec}^{\mathrm{R}}$ | BN18, This Study |
| *$bap1^{S551A}$* | *$vpvC^{\mathrm{W240R}}$, ΔrbmC, $bap1^{S551A}$, ΔVC1807::$P_{tac}$-mNeonGreen, $Spec^{\mathrm{R}}$* | Introduction of mutated *bap1* to Δ*bap1* Δ*rbmC* by cotransformation with Δ*VC1807*::*$P_{tac}$-mNeonGreen,* $\mathrm{Spec}^{\mathrm{R}}$ | BN19, This Study |
| *$bap1^{R118A}$* | *$vpvC^{\mathrm{W240R}}$, ΔrbmC, $bap1^{R118A}$, ΔVC1807::$P_{tac}$-mNeonGreen, $Spec^{\mathrm{R}}$* | Introduction of mutated *bap1* to Δ*bap1* Δ*rbmC* by cotransformation with Δ*VC1807*::*$P_{tac}$-mNeonGreen,* $\mathrm{Spec}^{\mathrm{R}}$ | BN20, This Study |
| *$bap1^{N550A}$* | *$vpvC^{\mathrm{W240R}}$, ΔrbmC, $bap1^{N550A}$, ΔVC1807::$P_{tac}$-mNeonGreen, $Spec^{\mathrm{R}}$* | Introduction of mutated *bap1* to Δ*bap1* Δ*rbmC* by cotransformation with Δ*VC1807*::*$P_{tac}$-mNeonGreen,* $\mathrm{Spec}^{\mathrm{R}}$ | BN21, This Study |
| *$bap1^{R118M}$* | *$vpvC^{\mathrm{W240R}}$, ΔrbmC, $bap1^{R118M}$, ΔVC1807::$P_{tac}$-mNeonGreen, $Spec^{\mathrm{R}}$* | Introduction of mutated *bap1* to Δ*bap1* Δ*rbmC* by cotransformation with Δ*VC1807*::*$P_{tac}$-mNeonGreen,* $\mathrm{Spec}^{\mathrm{R}}$ | BN27, This Study |
| *$bap1^{T597A}$* | *$vpvC^{\mathrm{W240R}}$, ΔrbmC, $bap1^{T597A}$, ΔVC1807::$P_{tac}$-mNeonGreen, $Spec^{\mathrm{R}}$* | Introduction of mutated *bap1* to Δ*bap1* Δ*rbmC* by cotransformation with Δ*VC1807*::*$P_{tac}$-mNeonGreen,* $\mathrm{Spec}^{\mathrm{R}}$ | JC01, This Study |

| | | | |
|---|---|---|---|
| *bap1*$^{Y88A}$ | *vpvC*$^{W240R}$, *ΔrbmC*, *bap1*$^{Y88A}$, *ΔVC1807::P*$_{tac}$*-mNeonGreen*, *Spec*$^{R}$ | Introduction of mutated *bap1* to *Δbap1 ΔrbmC* by cotransformation with *ΔVC1807::P*$_{tac}$*-mNeonGreen*, Spec$^{R}$ | JC02, This Study |
| **Strains Used for *in situ* Staining** | | | |
| *bap1-3XFLAG* | *vpvC*$^{W240R}$, *bap1-3XFLAG*, *ΔVC1807::P*$_{tac}$*-mNeonGreen*, Spec$^{R}$ | 3XFLAG tagged Bap1 | JY488[3] |
| *ΔrbmC* | *vpvC*$^{W240R}$, *ΔrbmC*, *ΔVC1807::P*$_{tac}$*-mNeonGreen*, Spec$^{R}$ | Non-tagged Bap1 with *rbmC* deleted | ZJ033, This Study |
| *ΔrbmC bap1-3XFLAG* | *vpvC*$^{W240R}$, *ΔrbmC*, *bap1-3XFLAG*, *ΔVC1807::P*$_{tac}$*-mNeonGreen*, Spec$^{R}$ | 3XFLAG tagged Bap1 with *rbmC* deleted | ZJ065, This Study |
| *bap1*$^{Y88F}$*-3XFLAG* | *vpvC*$^{W240R}$, *ΔrbmC*, *bap1*$^{Y88F}$*-3XFLAG*, *ΔVC1807::P*$_{tac}$*-mNeonGreen, Kan*$^{R}$ | 3XFLAG tagged Bap1$^{Y88F}$ with *rbmC* deleted | BN22, This Study |
| *bap1*$^{W164A}$*-3XFLAG* | *vpvC*$^{W240R}$, *ΔrbmC*, *bap1*$^{W164A}$*-3XFLAG*, *ΔVC1807::P*$_{tac}$*-mNeonGreen, Kan*$^{R}$ | 3XFLAG tagged Bap1$^{W164A}$ with *rbmC* deleted | BN23, This Study |
| *bap1*$^{Y88W}$*-3XFLAG* | *vpvC*$^{W240R}$, *ΔrbmC*, *bap1*$^{Y88W}$*-3XFLAG*, *ΔVC1807::P*$_{tac}$*-mNeonGreen, Kan*$^{R}$ | 3XFLAG tagged Bap1$^{Y88W}$ with *rbmC* deleted | BN24, This Study |
| *bap1*$^{K553A}$*-3XFLAG* | *vpvC*$^{W240R}$, *ΔrbmC*, *bap1*$^{K553A}$*-3XFLAG*, *ΔVC1807::P*$_{tac}$*-mNeonGreen, Kan*$^{R}$ | 3XFLAG tagged Bap1$^{K553A}$ with *rbmC* deleted | BN25, This Study |
| *bap1*$^{S551K}$*-3XFLAG* | *vpvC*$^{W240R}$, *ΔrbmC*, *bap1*$^{S551K}$*-3XFLAG*, *ΔVC1807::P*$_{tac}$*-mNeonGreen, Kan*$^{R}$ | 3XFLAG tagged Bap1$^{S551KA}$ with *rbmC* deleted | BN26, This Study |
| *bap1*$^{L598K}$*-3XFLAG* | *vpvC*$^{W240R}$, *ΔrbmC*, *bap1*$^{L598K}$*-3XFLAG*, *ΔVC1807::P*$_{tac}$*-mNeonGreen, Kan*$^{R}$ | 3XFLAG tagged Bap1$^{L598K}$ with *rbmC* deleted | BN28, This Study |
| *bap1*$^{S551A}$*-3XFLAG* | *vpvC*$^{W240R}$, *ΔrbmC*, *bap1*$^{S551A}$*-3XFLAG*, *ΔVC1807::P*$_{tac}$*-mNeonGreen, Kan*$^{R}$ | 3XFLAG tagged Bap1$^{S551A}$ with *rbmC* deleted | BN29, This Study |

| | | | |
|---|---|---|---|
| *bap1*$^{R118A}$*-3XFLAG* | *vpvC*$^{W240R}$, *ΔrbmC, bap1*$^{R118A}$*-3XFLAG, ΔVC1807::P*$_{tac}$*-mNeonGreen, Kan*$^{R}$ | 3XFLAG tagged Bap1$^{R118A}$ with *rbmC* deleted | BN30, This Study |
| *bap1*$^{N550A}$*-3XFLAG* | *vpvC*$^{W240R}$, *ΔrbmC, bap1*$^{N550A}$*-3XFLAG, ΔVC1807::P*$_{tac}$*-mNeonGreen, Kan*$^{R}$ | 3XFLAG tagged Bap1$^{N550A}$ with *rbmC* deleted | BN31, This Study |
| *bap1*$^{R118A, L598K}$*-3XFLAG* | *vpvC*$^{W240R}$, *ΔrbmC, bap1*$^{R118A, L598K}$*-3XFLAG, ΔVC1807::P*$_{tac}$*-mNeonGreen, Kan*$^{R}$ | 3XFLAG tagged Bap1$^{R118A, L598K}$ with *rbmC* deleted | BN32, This Study |
| *bap1*$^{R118M}$*-3XFLAG* | *vpvC*$^{W240R}$, *ΔrbmC, bap1*$^{R118M}$*-3XFLAG, ΔVC1807::P*$_{tac}$*-mNeonGreen, Kan*$^{R}$ | 3XFLAG tagged Bap1$^{R118M}$ with *rbmC* deleted | BN33, This Study |
| *bap1*$^{T597A}$*-3XFLAG* | *vpvC*$^{W240R}$, *ΔrbmC, bap1*$^{T597A}$*-3XFLAG, ΔVC1807::P*$_{tac}$*-mNeonGreen, Kan*$^{R}$ | 3XFLAG tagged Bap1$^{R118M}$ with *rbmC* deleted | JC05, This Study |
| *bap1*$^{Y88A}$*-3XFLAG* | *vpvC*$^{W240R}$, *ΔrbmC, bap1*$^{Y88A}$*-3XFLAG, ΔVC1807::P*$_{tac}$*-mNeonGreen, Kan*$^{R}$ | 3XFLAG tagged Bap1$^{R118M}$ with *rbmC* deleted | JC06, This Study |
| **Strains used for Complementation Assays** | | | |
| *bap1*$^{S551K}$ *+P*$_{BAD}$*-bap1* | *pEVS* (*P*$_{BAD}$*_bap1*, Kan$^{R}$), *vpvC*$^{W240R}$, *ΔrbmC, bap1*$^{S551K}$, *ΔVC1807::P*$_{tac}$*-mNeonGreen*, Spec$^{R}$ | *pEVS* containing *bap1* with an arabinose inducible promoter was mated into the indicated strain for complementation | BN34, This Study |
| *bap1*$^{L598K}$ *+P*$_{BAD}$*-bap1* | *pEVS* (*P*$_{BAD}$*_bap1*, Kan$^{R}$), *vpvC*$^{W240R}$, *ΔrbmC, bap1*$^{L598K}$, *ΔVC1807::P*$_{tac}$*-mNeonGreen*, Spec$^{R}$ | *pEVS* containing *bap1* with an arabinose inducible promoter was mated into the indicated strain for complementation | BN35, This Study |
| *bap1*$^{S551A}$ *+P*$_{BAD}$*-bap1* | *pEVS* (*P*$_{BAD}$*_bap1*, Kan$^{R}$), *vpvC*$^{W240R}$, *ΔrbmC, bap1*$^{S551A}$, *ΔVC1807::P*$_{tac}$*-mNeonGreen*, Spec$^{R}$ | *pEVS* containing *bap1* with an arabinose inducible promoter was mated into the indicated strain for complementation | BN36, This Study |
| *bap1*$^{N550A}$ *+P*$_{BAD}$*-bap1* | *pEVS* (*P*$_{BAD}$*_bap1*, Kan$^{R}$), *vpvC*$^{W240R}$, *ΔrbmC, bap1*$^{N550A}$, *ΔVC1807::P*$_{tac}$*-mNeonGreen*, Spec$^{R}$ | *pEVS* containing *bap1* with an arabinose inducible promoter was mated into the indicated strain for complementation | BN37, This Study |

| | | | |
|---|---|---|---|
| *bap1*$^{R118, L598K}$ +$P_{BAD}$-*bap1* | *pEVS* ($P_{BAD}$_*bap1*, Kan$^R$), *vpvC*$^{W240R}$, Δ*rbmC*, *bap1*$^{R118A, L598K}$, Δ*VC1807*::$P_{tac}$-*mNeonGreen*, Spec$^R$ | *pEVS* containing *bap1* with an arabinose inducible promoter was mated into the indicated strain for complementation | BN38, This Study |
| **Strains used for Rheological Measurement** | | | |
| Δ*pomA* | *vpvC*$^{W240R}$, Δ*rbmC*, Δ*pomA*, Δ*VC1807*::$P_{tac}$-*mNeonGreen, Kan*$^R$ | Deletion of *pomA* by cotransformation ZJ033 with Δ*VC1807*::$P_{tac}$-*mNeonGreen,* Kan$^R$ | JC15, This Study |
| Δ*pomA*, *bap1*$^{L598K}$ | *vpvC*$^{W240R}$, Δ*rbmC*, Δ*pomA*, *bap1*$^{L598K}$, Δ*VC1807*::$P_{tac}$-*mNeonGreen, Kan*$^R$ | Deletion of *pomA* by cotransformation BN18 with Δ*VC1807*::$P_{tac}$-*mNeonGreen,* Kan$^R$ | JC07, This Study |
| Δ*pomA*, *bap1*$^{S551K}$ | *vpvC*$^{W240R}$, Δ*rbmC*, Δ*pomA*, *bap1*$^{S551K}$, Δ*VC1807*::$P_{tac}$-*mNeonGreen, Kan*$^R$ | Deletion of *pomA* by cotransformation BN17 with Δ*VC1807*::$P_{tac}$-*mNeonGreen,* Kan$^R$ | JC08, This Study |
| Δ*pomA*, *bap1*$^{S551A}$ | *vpvC*$^{W240R}$, Δ*rbmC*, Δ*pomA*, *bap1*$^{S551A}$, Δ*VC1807*::$P_{tac}$-*mNeonGreen, Kan*$^R$ | Deletion of *pomA* by cotransformation BN19 with Δ*VC1807*::$P_{tac}$-*mNeonGreen,* Kan$^R$ | JC09, This Study |
| Δ*pomA*, *bap1*$^{R118A, L598K}$ | *vpvC*$^{W240R}$, Δ*rbmC*, Δ*pomA*, *bap1*$^{R118A, L598K}$, Δ*VC1807*::$P_{tac}$-*mNeonGreen, Kan*$^R$ | Deletion of *pomA* by cotransformation BN15 with Δ*VC1807*::$P_{tac}$-*mNeonGreen,* Kan$^R$ | JC10, This Study |

### SUPPLEMENTARY FIGURES

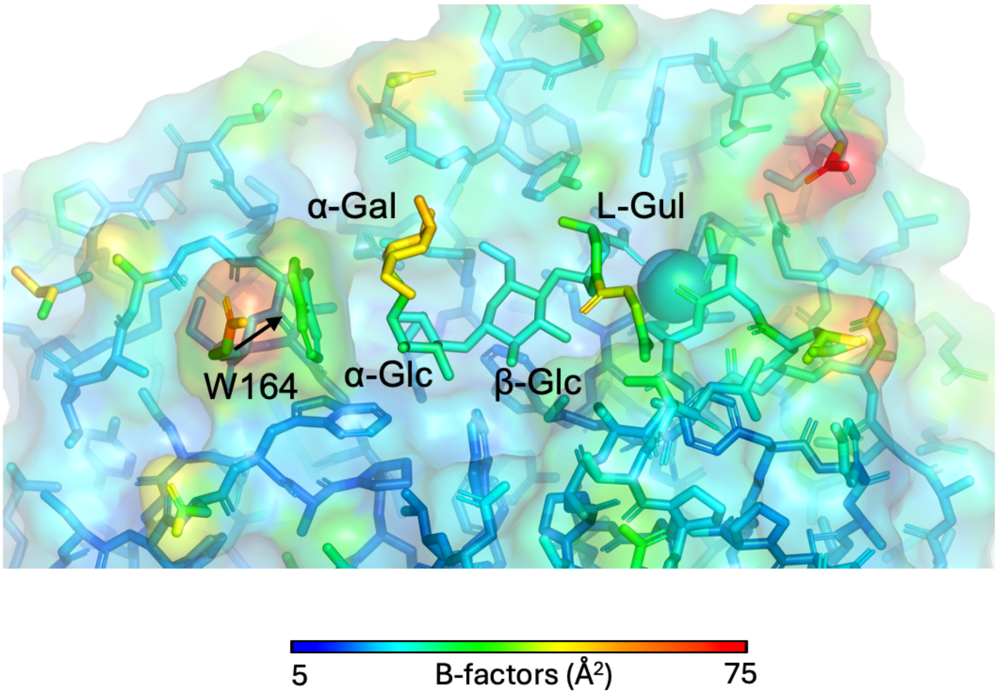

**Figure S1.** View of VPS binding site colored by crystallographic B-factor. The α-Gal moiety exhibits overall higher B-factors and less well-defined electron density than the rest of the VPS tetramer.

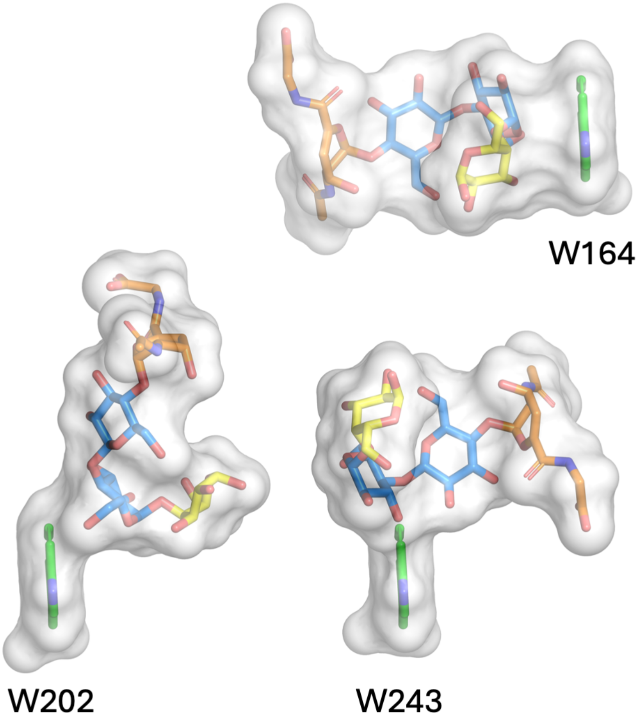

**Figure S2.** Stick/Space-filling representation of interactions between VPS and four adjacent tryptophan amino acid residues. To make this image, four copies of VPS with W164, W202, W243, and W599 individually represented were aligned on the tryptophan residue. VPS sugars are colored as in Fig. 1D.

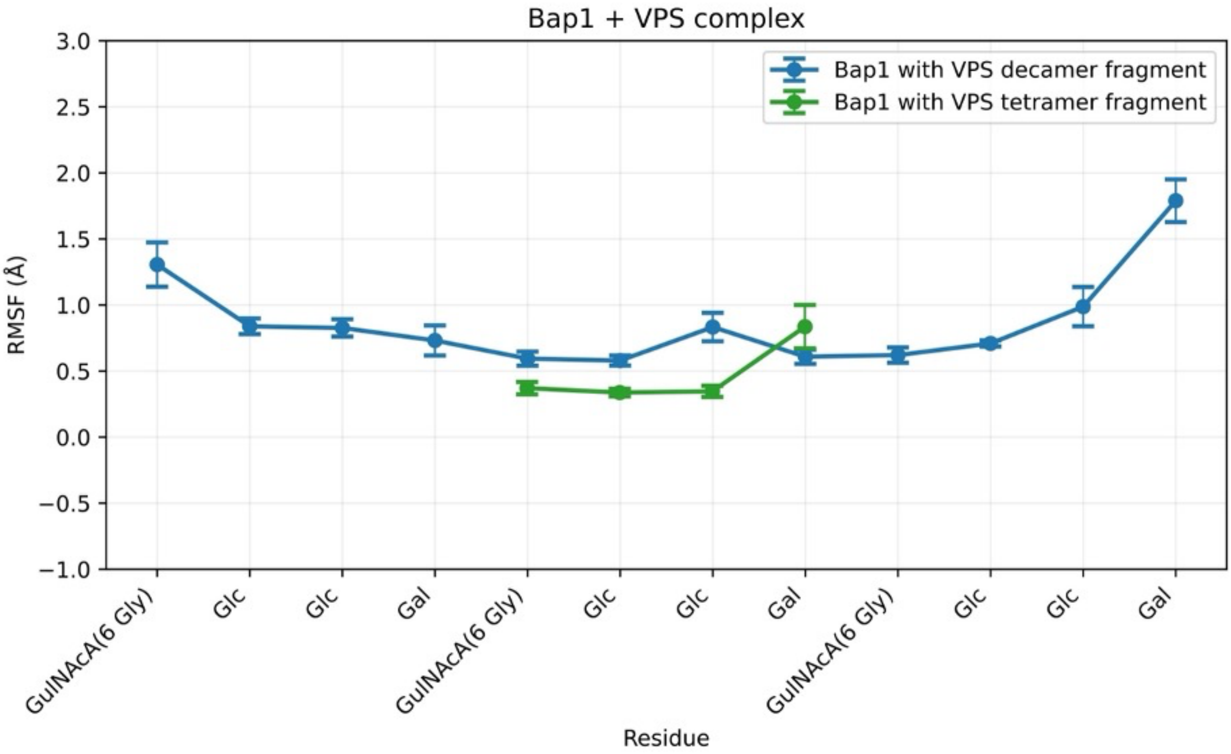

**Figure S3.** Root-mean-squared fluctuation (RMSF) analysis of the oligosaccharide (VPS) residues from MD simulations for Bap1 complexes, highlighting residue-level flexibility and dynamics.

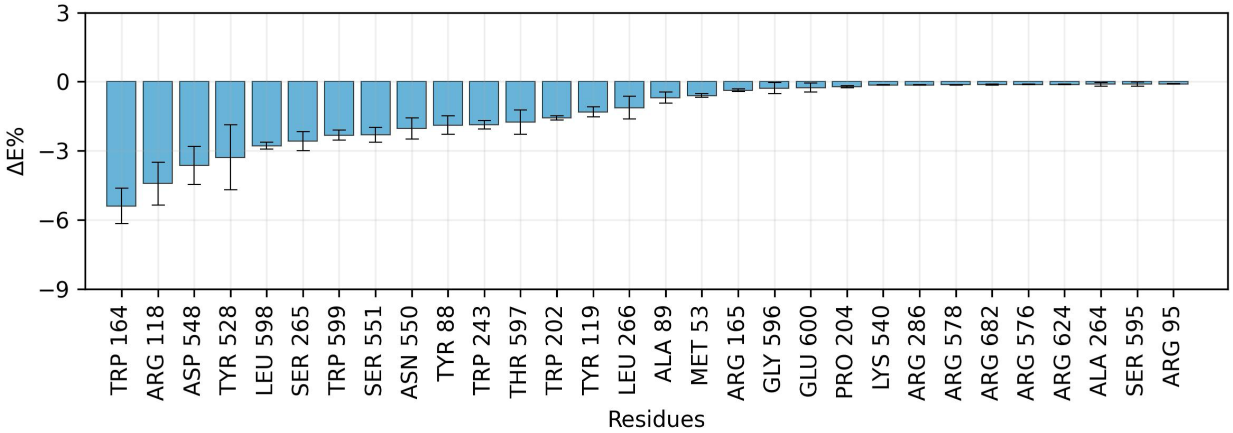

**Figure S4.** Per-residue energy decomposition for Bap1-VPS tetramer complex. The plot displays the percentage contribution of the top 30 residues to the total binding energy (ΔE). These residues are most critical to the formation and stability of the complex (Table S7).

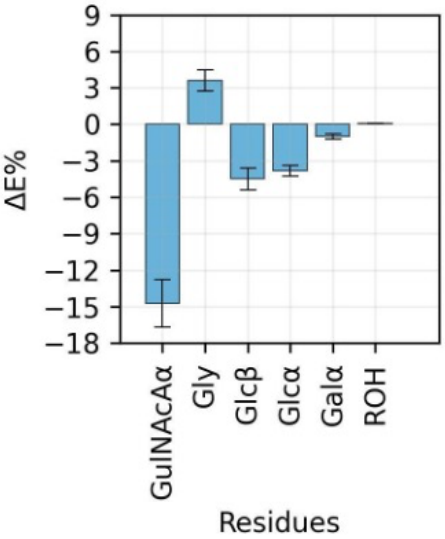

**Figure S5.** Percentage contribution to total interaction energy made by individual ligand residues in the Bap1 complexed with a tetrameric VPS fragment (data in Table S8).

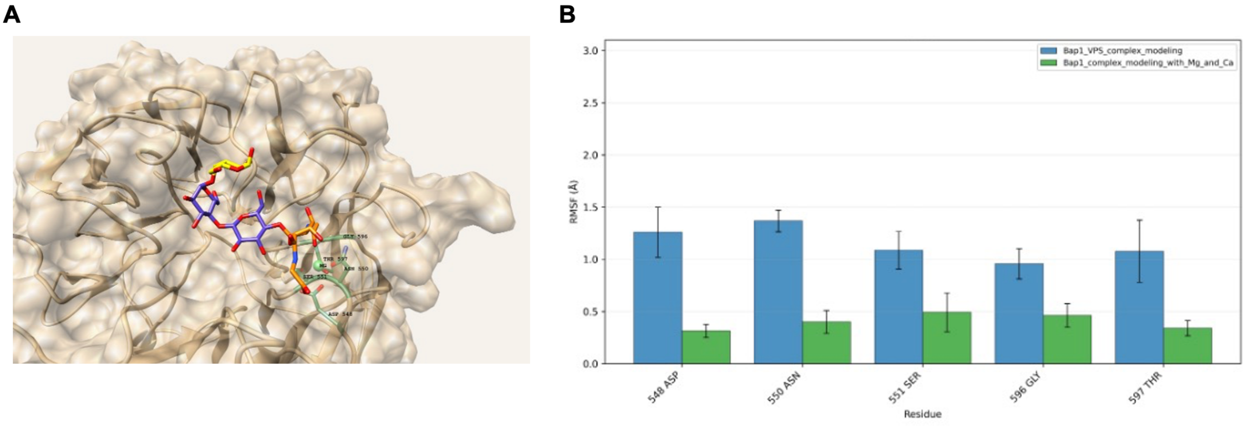

**Figure S6.** (A) Image and (B) RMSF of key Bap1 protein residues mediating $Mg^{2+}$ ion interactions in the Bap1-VPS co-complex as observed in the MD simulation. Bars represent MD simulations in the absence (blue) and presence (green) of $Mg^{2+}$ and $Ca^{2+}$ ions.

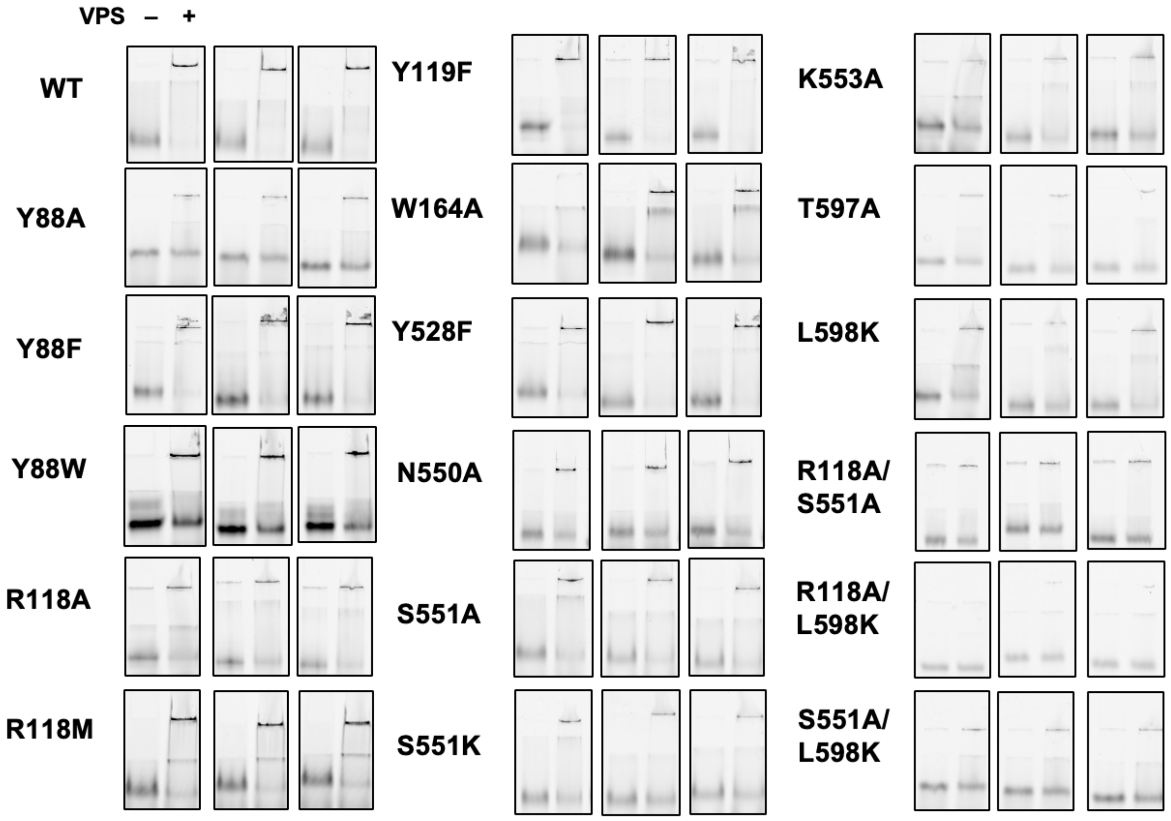

**Figure S7.** Individual fluorescence native gel images used to produce Fig. 4A. Bap1-GFP$_{UV}$ was imaged using a Typhoon fluorescence gel scanner as described in Materials and Methods.

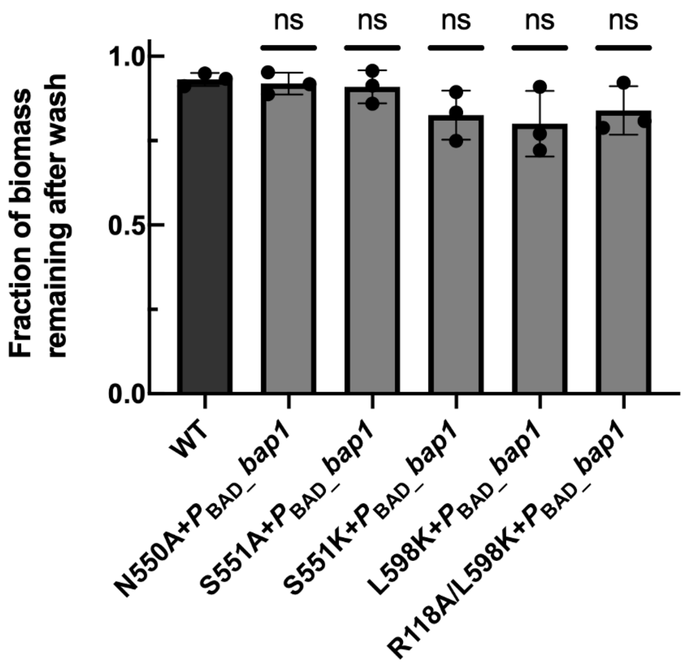

**Figure S8.** Biofilm adhesion assay showing that the biofilms with mutation leading to defective *bap1* can be complemented by expressing WT *bap1* induced by 0.2% arabinose on a plasmid. Statistical significance was determined using an unpaired, two-tailed *t*-test with Welch's correction compared to WT. ns = not significant.

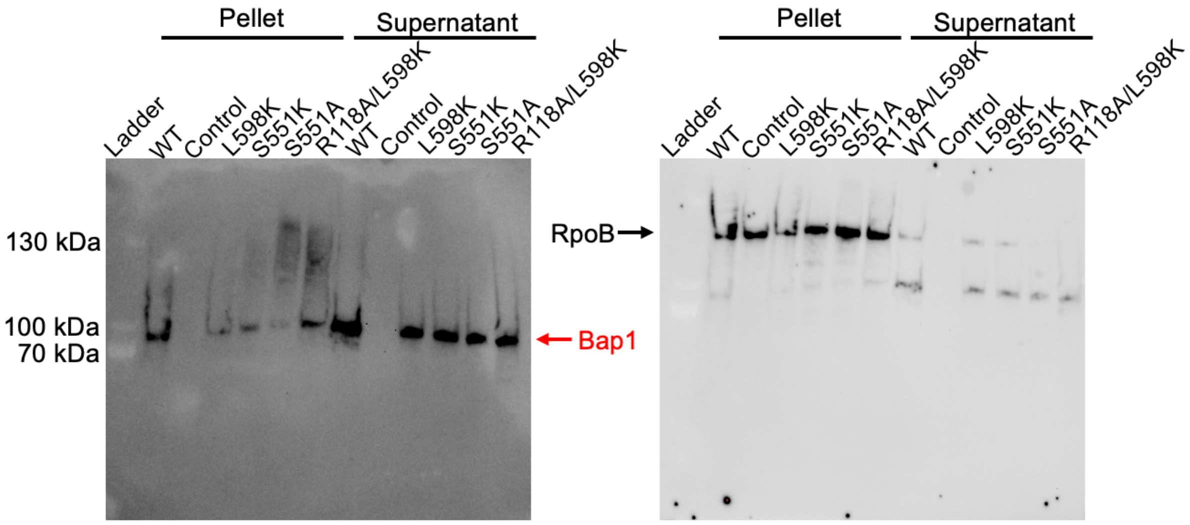

**Figure S9**. Western Blot showing normal production and secretion of defective Bap1 mutants in *V. cholerae*. A Δ*vpsL* background was used for all strains, to release Bap1 from the pellet into the supernatant. A strain without the 3×FLAG tag was used as a negative control. RpoB was used as a loading control as well as a control for cytosolically localized proteins.

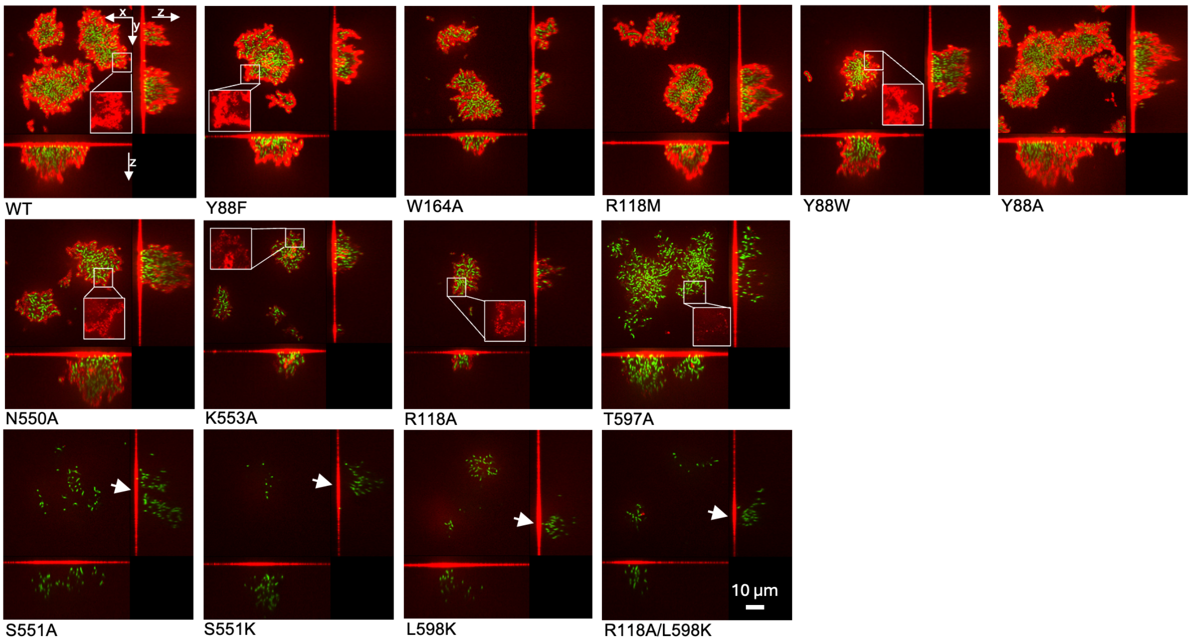

**Figure S10.** Cross-sectional confocal images of all *V. cholerae* mutants described in Fig. 4B,C, imaged using anti-FLAG antibody conjugated to Cy3 for spatial localization of Bap1 (red). *V. cholerae* cells constitutively express mNeonGreen (green). Boxes show zoomed-in features of the biofilm matrix. *z*-slices along the *xz* and *yz* axis are shown to the right and bottom of each image. Bap1 signals are observed in every case along the glass surface, particularly underneath the biofilm clusters, indicating successful secretion and adsorption to the glass substrata, regardless of whether the mutant Bap1 can bind to VPS and form envelopes surrounding biofilms clusters.

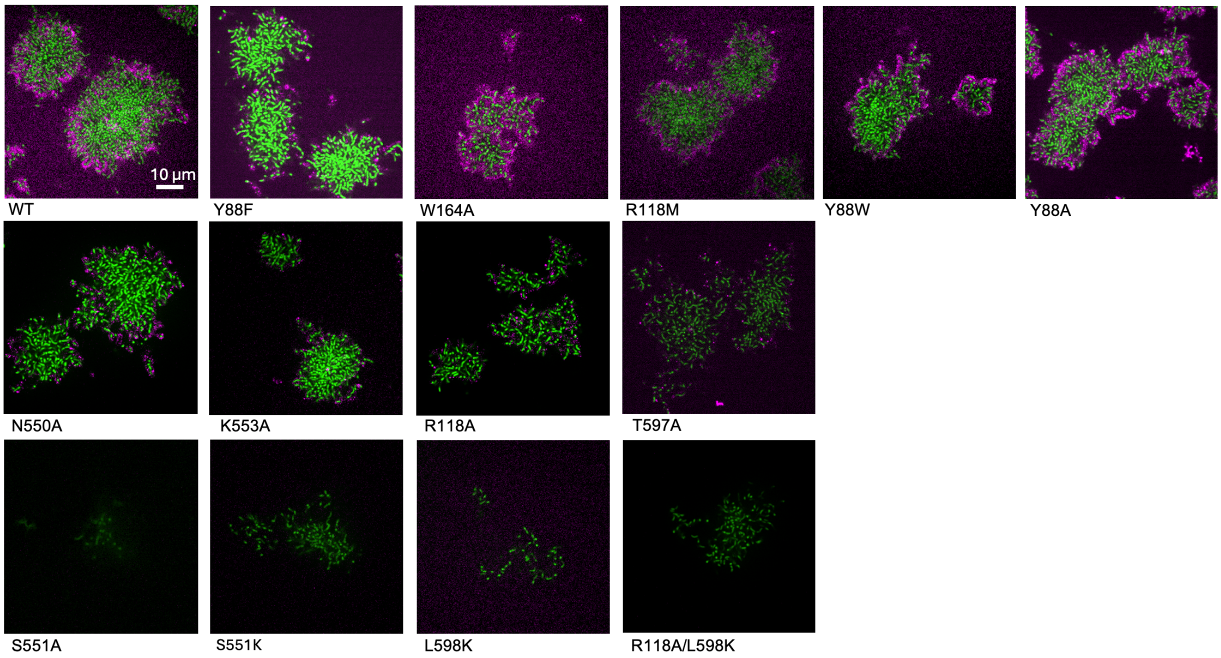

**Figure S11.** Cross-sectional confocal images of all *V. cholerae* mutants described in Fig. 4B,C, imaged using wheat germ agglutinin (WGA) conjugated to AlexaFluor647 for spatial localization of VPS (magenta). *V. cholerae* cells constitutively express mNeonGreen (green). Bap1 mutants are defective in binding VPS also shows reduced VPS signals. In previous work, it was shown that WGA staining signals are absent in the Δ*bap1*Δ*rbmC*, suggesting that at least one functional β-propeller is needed to organize VPS in a manner stainable by WGA.

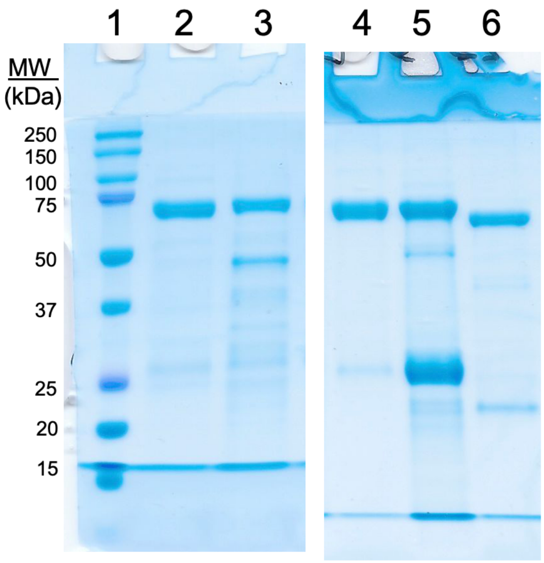

**Figure S12.** Purified proteins used in DLS experiments described in Fig. 5A. Lanes are as follows: Lane 1 – Molecular weight standards, Lane 2 – WT Bap1, Lane 3 – Bap1 R118A, Lane 4 – Bap1 L598K, Lane 5 – Bap1 R118A/L598K, Lane 6 – RbmB K227A.

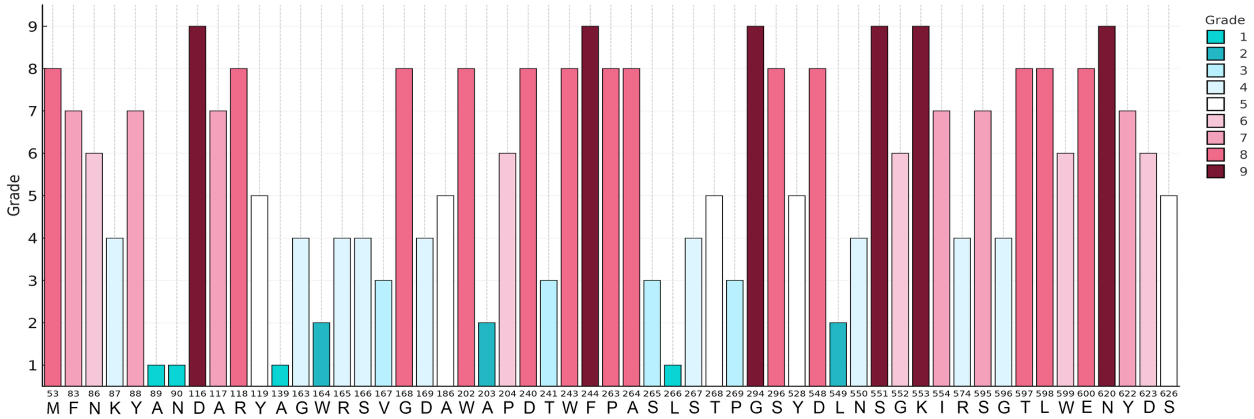

**Figure S13.** Sequence conservation of individual VPS-contacting Bap1 and RbmC amino acids. A higher grade signifies higher degree of conservation and coloring scheme matches Fig. 5C.

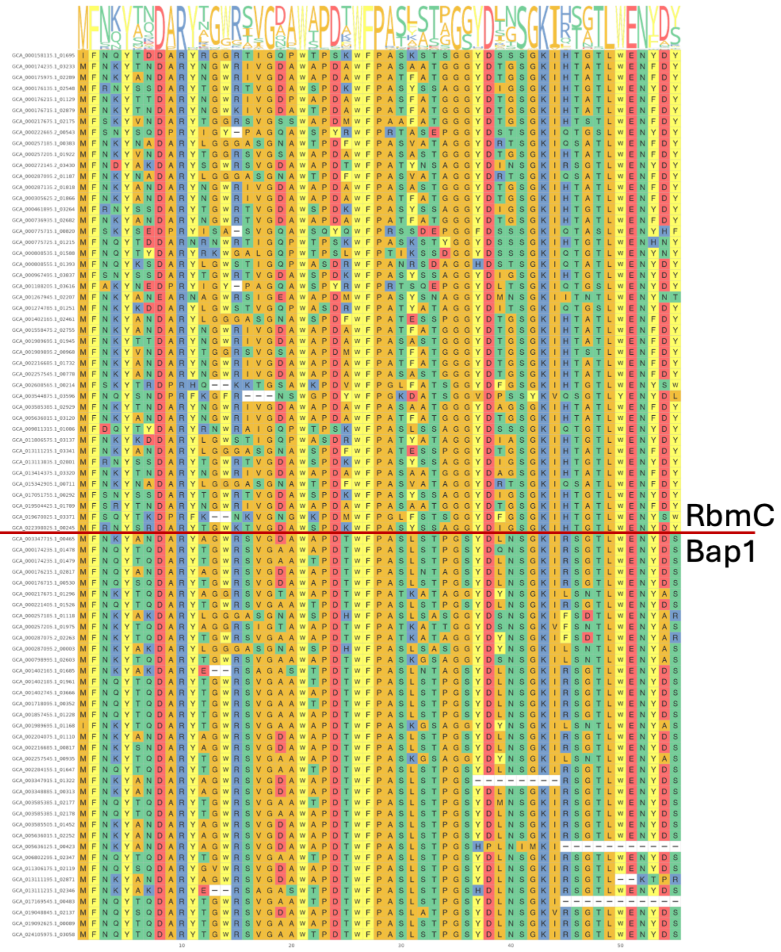

*V. cholerae* Bap1 residue

1. MET 53
2. PHE 83
3. ASN 86
4. LYS 87
5. TYR 88
6. ALA 89
7. ASN 90
8. ASP 116
9. ALA 117
10. ARG 118
11. TYR 119
12. ALA 139
13. GLY 163
14. TRP 164
15. ARG 165
16. SER 166
17. VAL 167
18. GLY 168
19. ASP 169
20. ALA 186
21. TRP 202
22. ALA 203
23. PRO 204
24. ASP 240
25. THR 241
26. TRP 243
27. PHE 244
28. PRO 263
29. ALA 264
30. SER 265
31. LEU 266
32. SER 267
33. THR 268
34. PRO 269
35. GLY 294
36. SER 296
37. TYR 528
38. ASP 548
39. LEU 549
40. ASN 550
41. SER 551
42. GLY 552
43. LYS 553
44. ILE 554
45. ARG 574
46. SER 595
47. GLY 596
48. THR 597
49. LEU 598
50. TRP 599
51. GLU 600
52. ASN 620
53. TYR 622
54. ASP 623
55. SER 626

**Figure S14.** Alignment of VPS-contacting RbmC and Bap1 amino acids showing conservation of individual amino acid positions. Residue numbering indicated in list on the left.

### Supplemental References

1. Yan, J., Nadell, C. D. & Bassler, B. L. Environmental fluctuation governs selection for plasticity in biofilm production. *ISME J.* **11**, 1569–1577 (2017).
2. Yan, J., Sharo, A. G., Stone, H. A., Wingreen, N. S. & Bassler, B. L. *Vibrio cholerae* biofilm growth program and architecture revealed by single-cell live imaging. *Proc. Natl. Acad. Sci. USA* **113**, e5337-5343 (2016).
3. Nijjer, J. *et al.* Mechanical forces drive a reorientation cascade leading to biofilm self-patterning. *Nat. Commun.* **12**, 6632 (2021).
4. Huang, X. *et al. Vibrio cholerae* biofilms use modular adhesins with glycan-targeting and nonspecific surface binding domains for colonization. *Nat Commun* **14**, 2104 (2023).
